# Shifting carbon flux from non-transient starch to lipid allows oil accumulation in transgenic tobacco leaves

**DOI:** 10.1101/2020.05.15.098632

**Authors:** Kevin L. Chu, Lauren M. Jenkins, Sally R. Bailey, Shrikaar Kambhampati, Somnath Koley, Kevin Foley, Jennifer J. Arp, Kirk J. Czymmek, Philip D. Bates, Doug K. Allen

**Author notes:** Author contributions: D.K.A. and P.D.B. conceived the original research plans; K.L.C., L.M.J., and S.R.B. performed the experiments, with assistance from S.Ka, K.F., and J.A.; K. L.C., L.M.J., S.Ka, S.Ko, K.F., and D.K.A. analyzed the data and contributed to scope; L. M.J. contributed expertise for the newly-developed acyl-ACP approach; K.J.C. performed all microscopy analysis; and K.L.C. and D.K.A. wrote the article with contribution from P.D.B. and was approved by all authors. The author responsible for distribution of materials integral to the findings presented in this article in accordance with the policy described in the Instructions for Authors (www.plantphysiol.org) is: Doug K. Allen.

## Abstract

Plant leaf biomass is composed predominantly of carbohydrate and protein with less than 5% dry weight allocated to lipid and less than 1% of total lipid in the form of triacylglycerols (TAGs). The combined overexpression of multiple genes involved in different aspects of TAG synthesis and stabilization can result in TAG accumulation to over 30% dry weight in tobacco leaves, presumably requiring many metabolic adjustments within plant cells. The metabolic consequences to the combined source and sink capacities of high oil accumulating transgenic tobacco leaves compared to wild-type were inspected across development and photoperiod by utilizing foliar biomass components and ^13^CO_2_ flux through central carbon intermediates. Lipid biosynthesis was investigated through assessment of acyl-acyl carrier protein (ACP) pools using a recently derived quantification method that was extended to accommodate isotopic labeling. Lipids accumulated stepwise over plant development in the high-oil leaves, with ^13^CO_2_-labeling studies confirming increased carbon flux to lipids. The large increase in lipid content was concurrent with a decrease in foliar starch, with limited contribution from non-sucrose soluble sugars, indicating a redirection of carbon from starch to lipids. Starch accumulated non-transiently with plant age in wild-type leaves, suggesting an inherent capacity for a developmentally-regulated carbon sink in tobacco leaves that may have enabled the programmed altered carbon partitioning to lipids in transgenics. These studies provide insight into the metabolic plasticity of dual source-sink leaves over development and may in part explain recent successful leaf lipid engineering efforts in tobacco.

**One sentence summary:** Engineering high oil accumulation in tobacco leaves is enabled by inherent source-sink plasticity associated with non-transient foliar starch accumulation over development.

## Introduction

Photosynthetically competent leaves are the primary organ for carbon assimilation, producing metabolites for local needs or exporting sugars and amino acids to other tissues. Initially, young leaves are heterotrophic and rely on nutrients from the existing seedling prior to photosynthetic competence. However, leaves act as carbon source tissues during the majority of plant development, with loss of sugar import capacity occurring during dicotyledonous leaf maturation via a basipetal wave from tip to base (Turgeon, 2006). Mature leaves provide assimilated carbon for sink tissues including shoots, roots, and newer developing leaves, eventually resulting in the formation of a canopy of photosynthetically competent leaves in many plants. Later in the life cycle, mature leaves provide resources for reproductive tissue development, with biomass in older leaves or leaves that are sufficiently shaded to prevent significant carbon assimilation being remobilized to contribute 30-70% of their carbon and nitrogen to developing seeds (Himelblau and Amasino, 2001; Kichey et al., 2007; Diaz et al., 2008; Li et al., 2017). The metabolic role of a leaf thus changes over its lifetime and canopy location and can be characterized by photosynthetic measurements of carbon source activities such as the production, turnover, and export of sugars.

Leaves convert atmospheric carbon dioxide to reduced organic intermediates of metabolism and storage reserves through the Calvin-Benson cycle (CBC) (Bassham et al., 1954). The CBC is the source of precursors for biomass components including starch, amino acids for protein, lipids, and nucleotides for DNA and RNA (Raines and Paul, 2006; Smith and Stitt, 2007; Stitt and Zeeman, 2012; McClain and Sharkey, 2019). CO_2_ is combined with ribulose bisphosphate by ribulose bisphosphate carboxylase/ oxygenase (Rubisco), producing two molecules of phosphoglyceric acid (PGA). PGA is converted to other CBC intermediates, used to generate starch or fatty acids in the chloroplast, or transported to the cytosol for sucrose production and export from the leaf. Starch and sucrose production take place in different spatial locations despite sharing many of the same hexose phosphate intermediates. However, the immediate nucleotide sugar phosphate precursors are produced in distinct cellular compartments. UDP-glucose is a substrate for sucrose production in the cytosol, and ADP-glucose is the glucosyl-donor for starch granule production in chloroplasts (Stitt and Quick, 1989).

Starch is synthesized in leaves during the day to act as a temporary carbon storage reserve that is remobilized at night for cellular respiration or conversion to sucrose to sustain nocturnal growth in sink tissues (Geiger and Servaites, 1994; Ludewig and Flügge, 2013). Starch turnover involves disruption of granule packing via phosphorylation of exposed glucans followed by breakdown of the polysaccharide polymers to maltose and glucose that are exported to the cytosol for further metabolism (Lu et al., 2005; Zeeman et al., 2010; Stitt and Zeeman, 2012; Smith and Zeeman, 2020). In *Arabidopsis thaliana*, 30-50% or more of assimilated carbon is converted to starch (Zeeman et al., 2010), depending on photoperiod length (Mengin et al., 2017). The regulation of starch levels is important to maximize carbon use efficiency and avoid carbon starvation during the night. In Arabidopsis, starch levels follow a diurnal pattern such that changing day and night lengths results in altered linear rates of starch production and turnover, with about 95% of the starch always being consumed by the end of the night (Lu et al., 2005; Smith and Stitt, 2007; Gibon et al., 2009; Graf et al., 2010; Graf and Smith, 2011; Mengin et al., 2017). This tight control over starch turnover may be relaxed during long-day photoperiods (Sulpice et al., 2014; Baerenfaller et al., 2015), with starch turnover ensuing alongside synthesis in the light to prevent overaccumulation and maintain optimal levels of sucrose for the start of the light-dark transition (Fernandez et al., 2017). Through this coordination, the production and levels of foliar sucrose are also stably controlled over most of the light/dark period (Durand et al., 2018).

While diurnal patterns of foliar starch accumulation and degradation similar to Arabidopsis have been observed in other plants including potato (*Solanum tuberosum*) (Lorenzen and Ewing, 1992) and sugar beet (*Beta vulgaris*) (Fondy and Geiger, 1982), differing starch trends have been observed in some plants. Unlike Arabidopsis, certain crops accumulate starch in their leaves over development. Leaves from older tobacco (*Nicotiana tabacum*), soybean (*Glycine max*), and *Lotus japonicus* plants contain much higher starch levels than leaves from younger plants (Matheson and Wheatley, 1962; Matheson and Wheatley, 1963; Häusler et al., 1998; Ölçer et al., 2001; Ainsworth et al., 2007; Vriet et al., 2010). Tobacco and *Lotus* starch is not completely turned over by the end of night, indicating developmental accumulation of non-transient foliar starch pools or delayed processing due to long photoperiods (Matt et al., 1998). In contrast to Arabidopsis starchless mutants that grow poorly under normal photoperiods (Caspar et al., 1985; Lin et al., 1988; Caspar et al., 1991), starchless mutants of *Nicotiana sylvestris* are normal except under short day photoperiods (Huber and Hanson, 1992; Geiger et al., 1995), suggesting that starch may be less crucial as a daily carbon source in some plants. Starchless mutants of pea (*Pisum sativum*) (Harrison et al., 1998) and *Lotus japonicus* (Vriet et al., 2010) also bear more phenotypic similarity to tobacco than to Arabidopsis, implying that starch metabolism is more complex across different plant species (Graf and Smith, 2011; Smith and Zeeman, 2020) than observed in model species studies. Such inconsistencies across species challenge our understanding of growth processes and complicate studies on photosynthetic operation and carbon partitioning in plants.

Triacylglycerols (TAG) are another important form of carbon reserve in plants, reaching over 80% dry weight (DW) in oil palm mesocarp and 20-50% DW in seeds of common oilseed crops (Weselake et al., 2009; Baud and Lepiniec, 2010; Bourgis et al., 2011). Most leaves, however, accumulate only 2-5% of their DW biomass as lipids, primarily in the form of galactolipids for photosynthetic membranes. Less than 1.5% of the total lipid in leaves is in the form of energy-dense TAGs (Koiwai et al., 1983; Yang and Ohlrogge, 2009; Li-Beisson et al., 2013; Allen et al., 2015), where they play important roles in membrane lipid homeostasis and temporarily store free fatty acids released during senescence and various abiotic stresses (Kaup et al., 2002; Fan et al., 2014; Fan et al., 2017; Arisz et al., 2018; Ischebeck et al., 2020). Efforts to engineer increased production and accumulation of TAG in vegetative tissues for more energy-dense biomass have achieved the most success through the combined targeting of multiple aspects of developmental central carbon metabolism, TAG biosynthesis, and lipid droplet stabilization (the ‘Push’-‘Pull’-‘Package’-‘Protect’ approach) (Durrett et al., 2008; Slocombe et al., 2009; Vanhercke et al., 2014a; Weselake, 2016; Xu and Shanklin, 2016; Beechey-Gradwell et al., 2019; Vanhercke et al., 2019b; Parajuli et al., 2020). Recently, tobacco lines were reported with dramatically altered carbon partitioning to TAG in leaves (Andrianov et al., 2010; Vanhercke et al., 2014b; Vanhercke et al., 2017; Cai et al., 2019; Zhou et al., 2020). Some of these engineered tobacco lines produced over 30% TAG (DW) in leaves without drastic consequences on plant growth. These foliar TAG levels are the highest reported thus far in any engineered plant species, even when similar gene combinations are manipulated (reviewed in Vanhercke et al., 2019b). However, it is unclear how tobacco leaf metabolism is able to adapt to the increased demand for assimilated carbon incorporation into chloroplastic fatty acid synthesis.

The current study comparatively assesses carbon partitioning over development in wild-type (WT) tobacco and a high lipid-producing transgenic line overexpressing the *WR1, DGAT1, OLEOSIN*, and *LEC2* genes (hereafter referred to as the HO-LEC2 line) to address this question. Lipid, starch, and soluble sugar levels were characterized in wild type and HO-LEC2 transgenic leaves, with measurements at different times in the photoperiod across development. HO-LEC2 leaves accumulated more lipid than WT throughout development, starting at a young age. Lipid as a percentage of total biomass, however, did not change significantly in the HO-LEC2 line until after eight weeks of age when a rapid increase in lipid accumulation to over 30% DW occurred. One striking observation was that WT leaves accumulated a large pool of non-transient starch with development, whereas high lipid-producing HO-LEC2 leaves showed a significant drop in all foliar starch coinciding with the step increase in lipids at nine weeks. The findings imply a tradeoff in carbon between starch and lipids that was further scrutinized using ^13^CO_2_-labeling experiments. Dynamic fluxes through photosynthetic and carbohydrate metabolism as well as fatty acid synthesis were measured over several hours to gauge the production and turnover of starch, sucrose, and fatty acids in HO-LEC2 leaves against WT. Lipid biosynthesis was assessed using a recently developed method for quantifying acyl-ACP intermediates (Nam et al., 2020), modified to monitor ^13^C-label incorporation. Acyl-ACP quantitation also indicated diurnal changes in fatty acid synthesis comparable to starch. Our results reveal major changes in central carbon metabolism that occur during the vegetative stage in oil accumulating leaves and suggest why foliar lipid engineering efforts have been more successful in tobacco compared to other plant species.

## Results

### Lipid levels increase over development in a stepwise manner

The top expanded leaves of vegetative WT and HO-LEC2 transgenic tobacco plants in the greenhouse were sampled at midday at 26, 41, 55, 62, and 72 days after sowing (DAS) for developmental analysis of lipid production and accumulation. HO-LEC2 plants showed delayed seedling establishment and a 1/6^th^ reduction in leaf size compared with WT plants but reached comparable heights by 55 DAS and flowered at similar times. Neither line started flowering or bolting until after 77 DAS. Older HO-LEC2 leaves displayed increased leaf curling and developed a mottled yellow appearance that intensified with plant age, likely due to reduced chlorophyll levels (Fig.1A). HO-LEC2 leaves showed significantly higher lipid levels than WT at all ages tested, with WT leaves not exhibiting any major changes in lipid content over development (Fig. 1B). The elevated lipid content in HO-LEC2 was apparent early in development but did not change further until between 55 and 62 DAS when a dramatic increase (almost doubling) occurred to near final levels. The total lipid levels at 62 DAS are close to the maximum TAG levels previously reported (Vanhercke et al., 2017) in senescing leaves at seed setting, with large increases in the absolute amounts of palmitic, oleic, and linoleic fatty acid species in HO-LEC2 leaves (Supplemental Fig.S1) also closely matching previously reported results. No major differences in lipid levels and composition were detected between dawn and dusk for both WT and HO-LEC2 leaves at 62 DAS, suggesting little diurnal lipid turnover in either line (Supplemental Fig.S1). Confocal microscopy of 67 DAS leaves stained with BODIPY 505/515 showed large amounts of lipid droplets in their mesophyll cells, especially the palisade layer, that were not present in WT leaf cells (Fig.1C-D).

**Figure 1.**
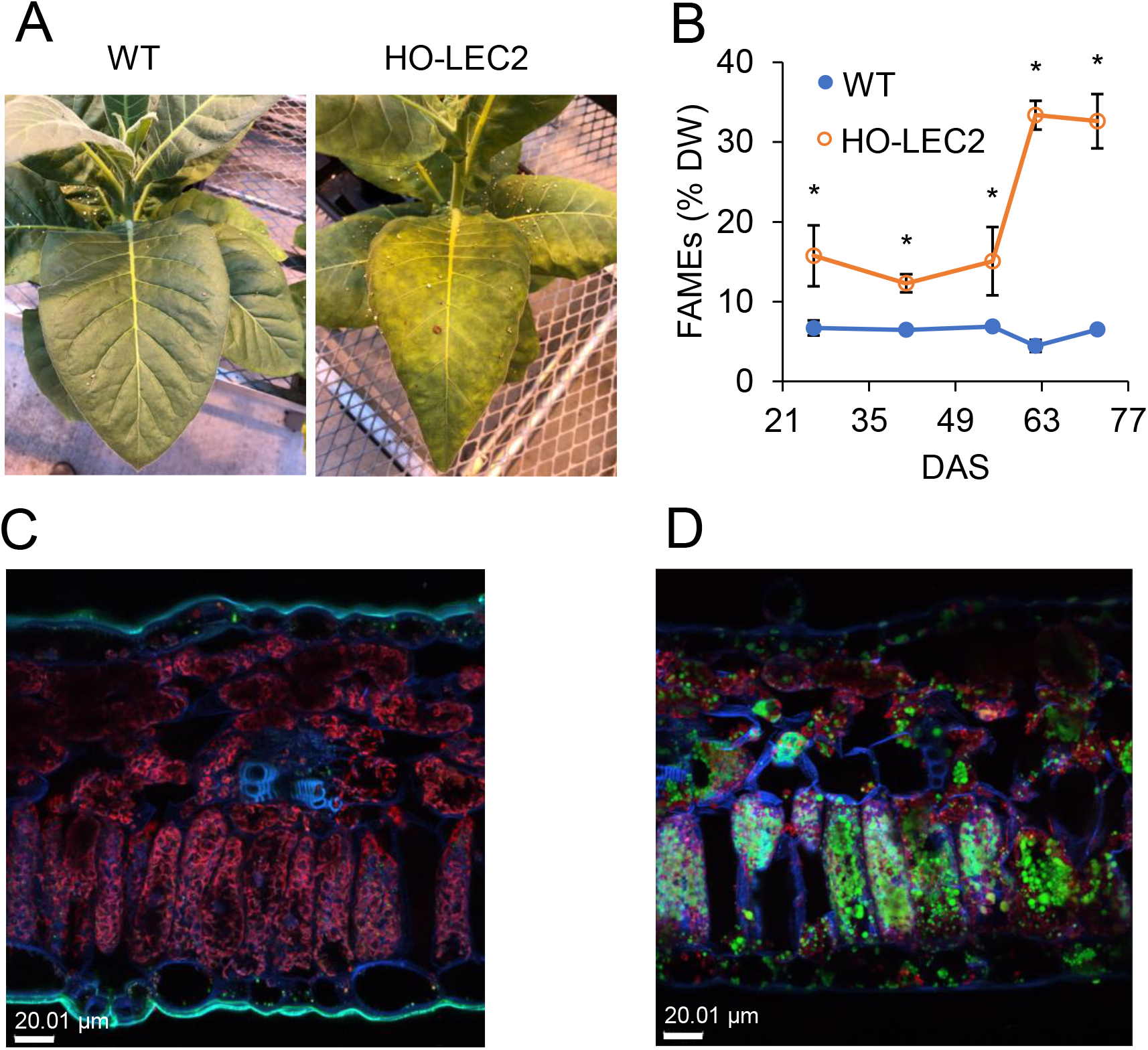
Acyl-lipid levels in WT and HO-LEC2 tobacco lines over development. A, The top expanded leaves of WT and HO-LEC2 plants at 63 DAS. B, Total fatty acid methyl ester (FAME) content of the top expanded leaf of WT and HO-LEC2 plants sampled at midday across development (DAS – days after sowing). The results given are mean ± SD of three to four biological replicates. Asterisks indicate significant differences from WT in the HO-LEC2 line (Student’s *t*-test, *P* < 0.05). C and D, Accumulation and distribution of lipid droplets in mesophyll cells stained with BODIPY 505/515 (green) in WT and HO-LEC2 leaf cross-sections. Confocal images are of merged BODIPY 505/515 emission, chlorophyll autofluorescence (magenta), and cell wall autofluorescence (cyan). Scale bars are 20.01 μm.

### Acyl-ACP measurements indicate enhanced production of fatty acids in HO-LEC2 leaves

Acyl-ACPs are common intermediates in the synthesis and elongation of fatty acids, and their measurement allows for an assessment of lipid production rates in addition to accumulation. The dynamic production of fatty acids for lipids was inspected in 63 DAS WT and HO-LEC2 leaves over the photoperiod by capitalizing on a recently developed technique to sensitively quantify acyl-ACPs of varying acyl chain lengths (Nam et al., 2020). Acyl-ACPs with acyl chain lengths from 2-16 carbons were consistently quantifiably in both WT and HO-LEC2 leaves. The levels of most acyl-ACPs were higher in HO-LEC2 than in WT, including a terminal C16:0-ACP product (Fig.2). The levels of acyl-ACPs in HO-LEC2 leaves also displayed diurnal fluctuations denoting elevated fatty acid synthesis in the light compared to the night. The lack of diurnal fluctuations in WT acyl-ACP levels underscore the decreased demand for lipids in these plants. These findings provide further evidence of enhanced lipid production in 63 DAS HO-LEC2 leaves consistent with the FAME accumulation data. Since both biomass accumulation and acyl-ACP measurements indicated a large increase in HO-LEC2 foliar lipid production by the nine-week age, plants at this stage were utilized for subsequent ^13^CO_2_-labeling studies.

**Figure 2.**
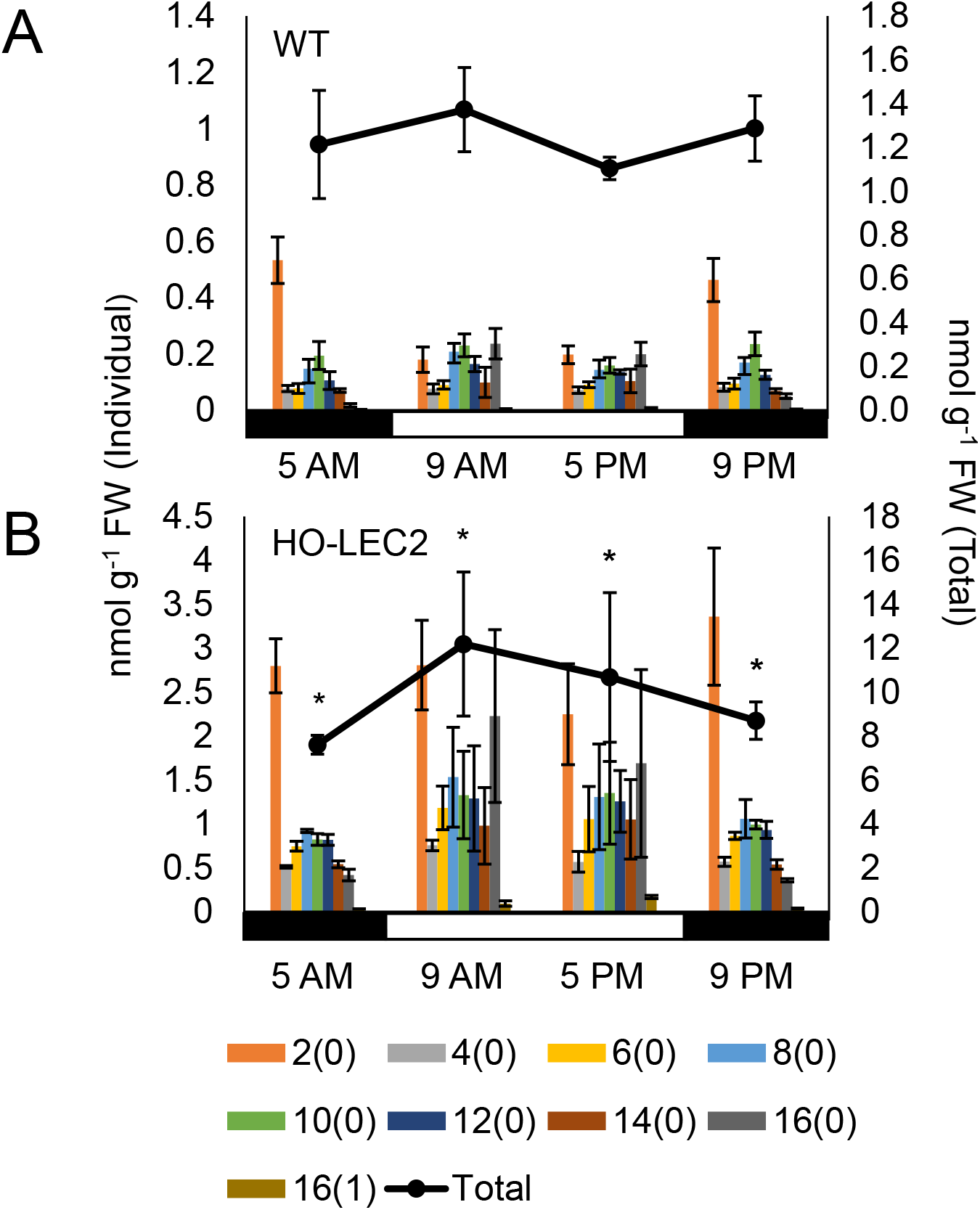
Diurnal levels of acyl-ACPs. Acyl-ACPs from the top expanded leaves of A, 63 DAS WT and B, HO-LEC2 plants were assessed over a 12 h L: 12 h D photoperiod. Individual ACPs are presented as bars from two-carbon to a terminal 16-carbon acyl-ACPs. Total acyl-ACP pools are indicated by the line and dot plot above. The results given are mean ± SD of three biological replicates. Asterisks indicate significant differences from WT of total acyl-ACP in the HO-LEC2 line (Student’s *t*-test, *P* < 0.05).

### WT but not HO-LEC2 leaves accumulate non-transient starch over development

The top expanded leaves of WT and HO-LEC2 plants were sampled for starch and soluble sugars over a 12-hour photoperiod at different developmental ages to investigate how increased leaf oil production affects carbon partitioning into carbohydrates. In WT leaves, foliar starch levels as a percentage of dry weight accumulated with plant age on top of diurnal fluctuations (Fig.3A-D). Interestingly, the amount of starch remaining at the end of night increased from 1.6% of DW at 41 DAS to 28.1% by 62 DAS before dropping to 19.2% at 72 DAS. Since foliar starch turnover occurs in the dark, the accumulated starch remaining at dawn is non-transient. Diurnal variations in starch, prominent in younger plants, became less pronounced with plant age due to starch accumulation. Leaf starch levels and diurnal patterns in the HO-LEC2 transgenic plants were initially slightly lower but similar to WT in younger plants. By 62 DAS, however, there was a significant drop in HO-LEC2 foliar starch at all timepoints to levels at or below those measured at 41 DAS (Fig.3). Diurnal starch patterns, while still apparent, also showed a noticeable decrease in amplitude. The drop in foliar starch levels is roughly concurrent with the significant accumulation of lipids in HO-LEC2 leaves (Fig.1A), suggesting a change in foliar carbon sink production. An acyl-sugar extraction step added at the start of a subset of starch assays did not alter measured starch levels, indicating that acyl-sugars did not erroneously contribute to starch measurements despite significant accumulation of acyl-sugars in Solanaceae leaves (Fobes et al., 1985; Severson et al., 1985).

**Figure 3.**
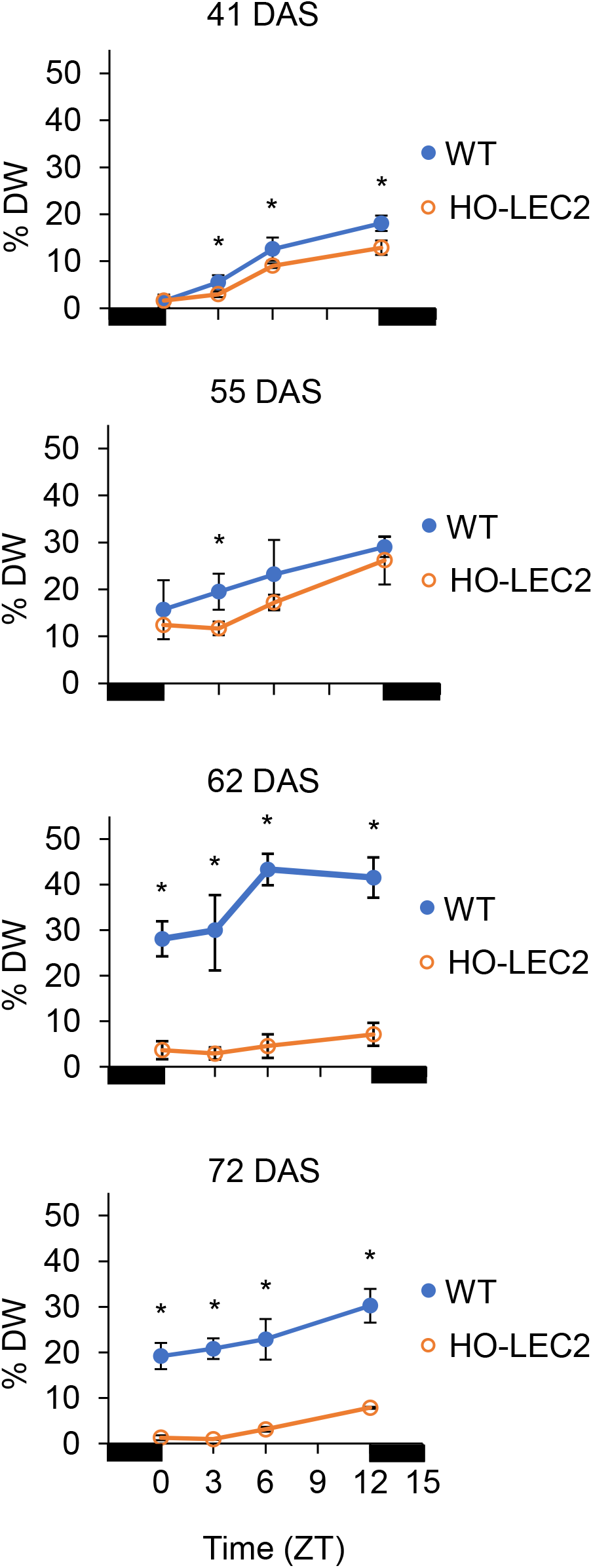
Diurnal starch levels in *WT* and HO-LEC2 leaves over development. Starch levels in the top expanded leaves of WT and HO-LEC2 over a 12 h L: 12 h D photoperiod at 41, 55, 62, and 72 DAS. Black bars represent darkness. The results given are mean ± SD of three to four biological replicates. Asterisks indicate significant differences from WT in the HO-LEC2 line (Student’s *t*-test, *P* < 0.05).

To investigate nocturnal starch turnover, nighttime levels of the disaccharide maltose were measured due to the unique position of maltose as an intermediate of starch-degradation not commonly found elsewhere in metabolism (Niittylä et al., 2004; Weise et al., 2004). WT and HO-LEC2 leaves were sampled at midnight every other day from 56 DAS to 64 DAS, the period spanning the large changes in lipid and starch levels. No significant changes in maltose levels were detected in HO-LEC2 leaves across this period (Fig.4A), indicating that increased rates of nighttime starch degradation are unlikely to be the reason for the decrease in HO-LEC2 starch levels. Midnight levels of maltose were found to be lower in HO-LEC2 leaves compared to WT, indicating either lower overall rates of starch turnover or increased turnover of maltose itself due to increased metabolic demands. These observations are consistent with the generally lower diurnal starch patterns measured in HO-LEC2 leaves.

**Figure 4.**
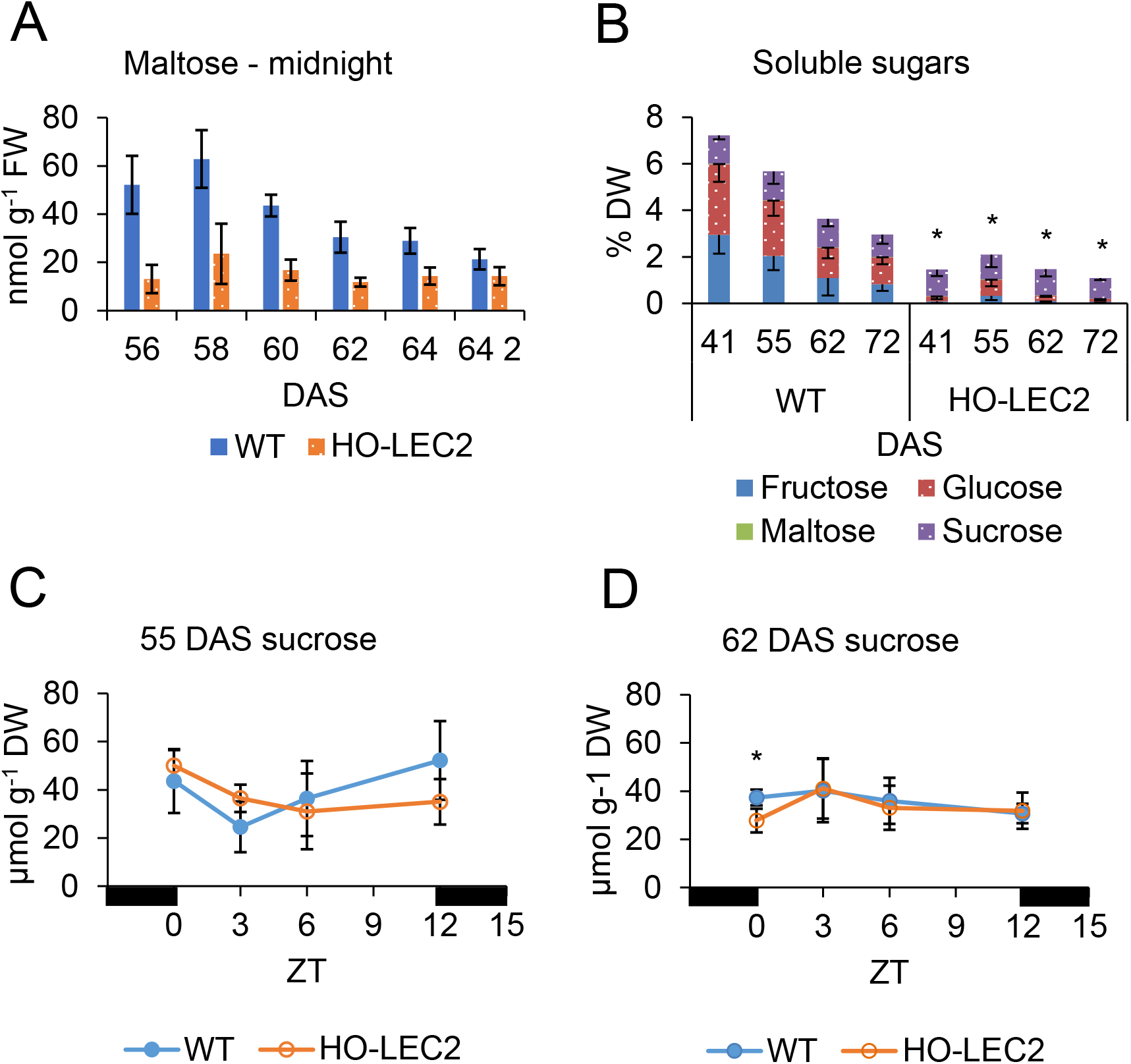
Soluble sugar changes in WT and HO-LEC2 leaves. A, Maltose levels of WT and HO-LEC2 leaves between 56 and 64 DAS at midnight. No statistically significant differences were found among HO-LEC2 levels using ANOVA. B, Soluble sugar levels in the top expanded leaf of WT and HO-LEC2 tobacco plants at midday at 41, 55, 62, and 72 DAS. Asterisks indicate significant differences from WT in total soluble sugars in the HO-LEC2 line (Student’s *t*-test, *P* < 0.05). C and D, Sucrose levels across the photoperiod of 55 and 62 DAS WT and HO-LEC2 leaves. The results given are mean ± SD of three to four biological replicates. Asterisks indicate significant differences from WT in the HO-LEC2 line (Student’s *t*-test, *P* < 0.05). No statistically significant differences were found over either photoperiod or development at midday for either WT or HO-LEC2 using ANOVA.

### Levels of non-sucrose soluble sugars are reduced in HO-LEC2 leaves

Several studies seeking to increase foliar carbon flux to lipids by targeting starch biosynthetic genes found a corresponding increase in soluble sugar levels along with lipids (Sanjaya et al., 2011; Yu et al., 2018; Xu et al., 2019). In HO-LEC2 leaves, however, levels of total soluble sugars at midday were significantly lower compared to WT across development, suggesting that starch biosynthetic enzymes do not appear to be overly compromised in HO-LEC2. The difference became less pronounced over development as WT sugar levels declined with plant age (Fig.4B). Both the developmental decline in WT sugars and the overall lower sugar levels in HO-LEC2 leaves were primarily due to decreased amounts of free fructose and glucose, a trend also apparent across the photoperiod (Supplemental Fig.S2). Sucrose levels in WT and HO-LEC2 leaves were generally not significantly different from each other over both photoperiod and development (Fig.4C-D) and did not exhibit any trends. The similarity of HO-LEC2 sucrose levels to WT across development suggests that the potential source strength of HO-LEC2 leaves during the day was not markedly affected by their increased lipid production.

### HO-LEC2 trades non-transient starch accumulation with lipid

The characterization of leaf biomass changes over development indicated that HO-LEC2 leaves accumulated acyl-lipids primarily at the expense of starch. Neither WT nor HO-LEC2 leaves exhibited any notable differences or trends in protein levels across development (Fig.5A), indicating that starch and lipid are the principal changing biomass components. The starch in WT leaves that accumulated over development is lost in HO-LEC2 leaves that instead store significant pools of lipid, thus preventing disruption of other biomass components. The biomass fractions of TAG and starch as carbon were calculated utilizing a 77-78% carbon composition for TAG based on the chemical formula of TAG molecules proportionally incorporating measured fatty acids (Supplemental Fig.S1) and a 43% carbon composition for starch using the chemical formula for maltotriose. From the calculations, 76% of the carbon from increased lipids can be accounted for by the difference in starch between WT and HO-LEC2 leaves at 62 DAS (Fig.5B). The availability of a large pre-existing non-transient carbon reserve in foliar plastids of older tobacco plants thus offers an explanation as to how high oil-accumulating HO-LEC2 leaves can produce large amounts of lipids without drastic penalties to plant growth and viability.

**Figure 5.**
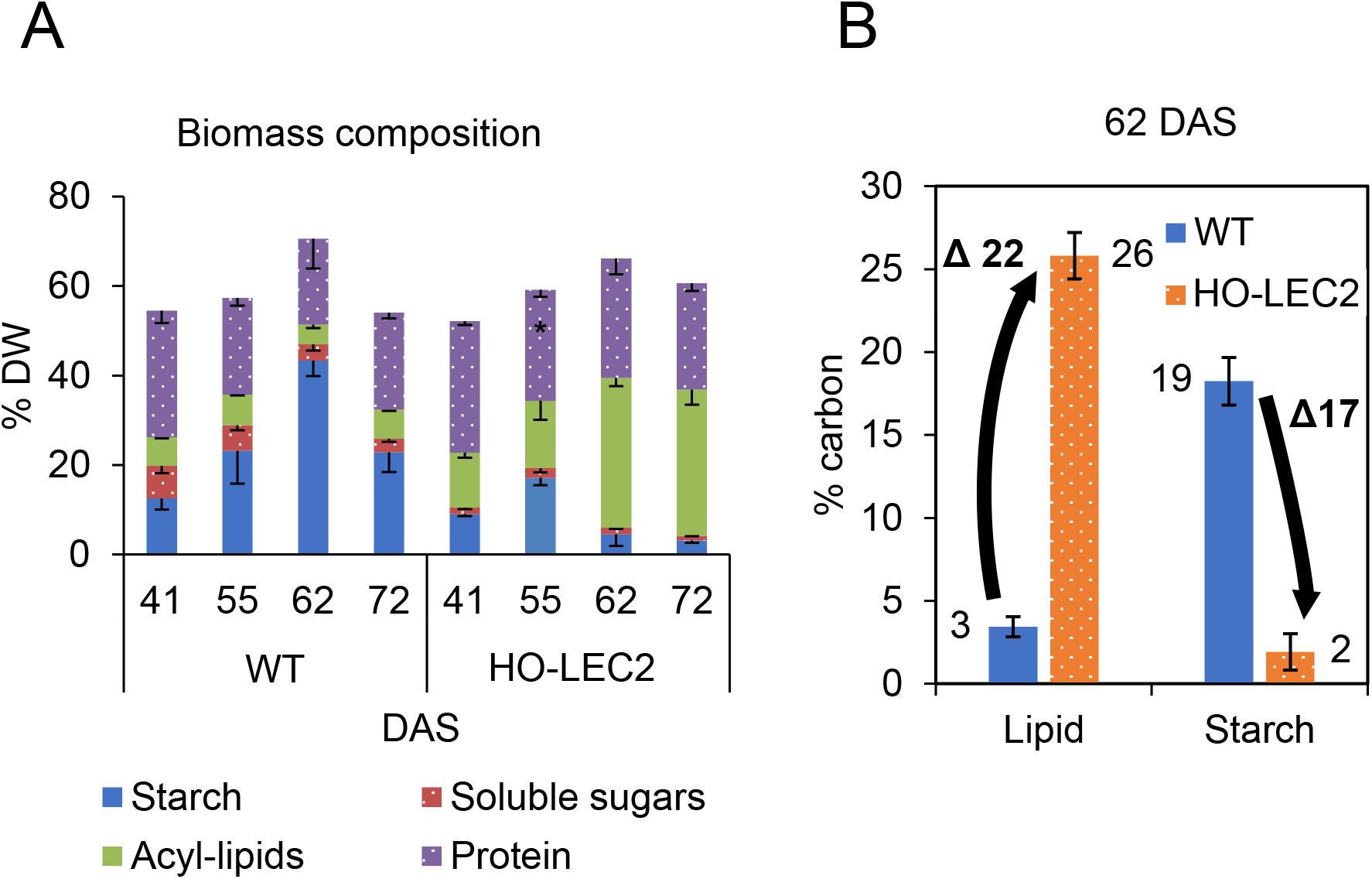
Biomass composition changes. A, Biomass components of the top expanded leaf of WT and HO-LEC2 tobacco plants over a 12 h L: 12 h D photoperiod at 41, 55, 62, and 72 DAS. The results given are mean ± SD of three to four biological replicates. Asterisks indicate significant differences from WT of protein in the HO-LEC2 line (Student’s *t*-test, *P* < 0.05). B, Percent carbon differences from WT and HO-LEC2 lipid and starch values at 62 DAS. Lipid percent carbon was calculated using the proportions and formulas of measured fatty acids, and starch percent carbon was calculated using the formula for maltotriose.

### HO-LEC2 leaves show reduced photosynthetic fluxes and starch production

Transient labeling of WT and HO-LEC2 leaves from nine-week old plants with a ^13^CO_2_ gas mixture was performed to assess photosynthetic carbon fluxes through various primary metabolic pathways. Total starch accumulation during five hour labeling experiments was compared with ^13^C-labeled starch accumulation as described in prior similar investigations (Fernandez et al., 2017). WT leaves synthesized both more total starch and more ^13^C-labeled starch than HO-LEC2 over the labeling period (Fig.6A-D), consistent with HO-LEC2 leaves allocating more carbon to produce lipids instead of starch. The smaller total starch pool size of HO-LEC2 plants relative to WT is consistent with the higher ^13^C-enrichment of starch glucose in HO-LEC2 (Fig.6B) and the lower amount of ^13^C-starch production observed (Fig.6C-D) in this line, since ^13^C-enrichment for a given flux will be higher for a smaller pool size. Labeled starch production rates were lower than total starch production rates for both WT and HO-LEC2 leaves (Fig.6D), indicating that significant starch turnover simultaneous with synthesis in the light as previously described in Arabidopsis (Fernandez et al., 2017) cannot explain the lower starch levels in HO-LEC2 leaves. If daytime starch turnover was the predominant explanation, the amount of ^13^C-starch produced over the labeling period should exceed or at least match the measured total starch production. The decrease in HO-LEC2 foliar starch levels thus do not appear to be due to elevated rates of either daytime or nighttime starch turnover and may instead result from altered starch synthesis.

**Figure 6.**
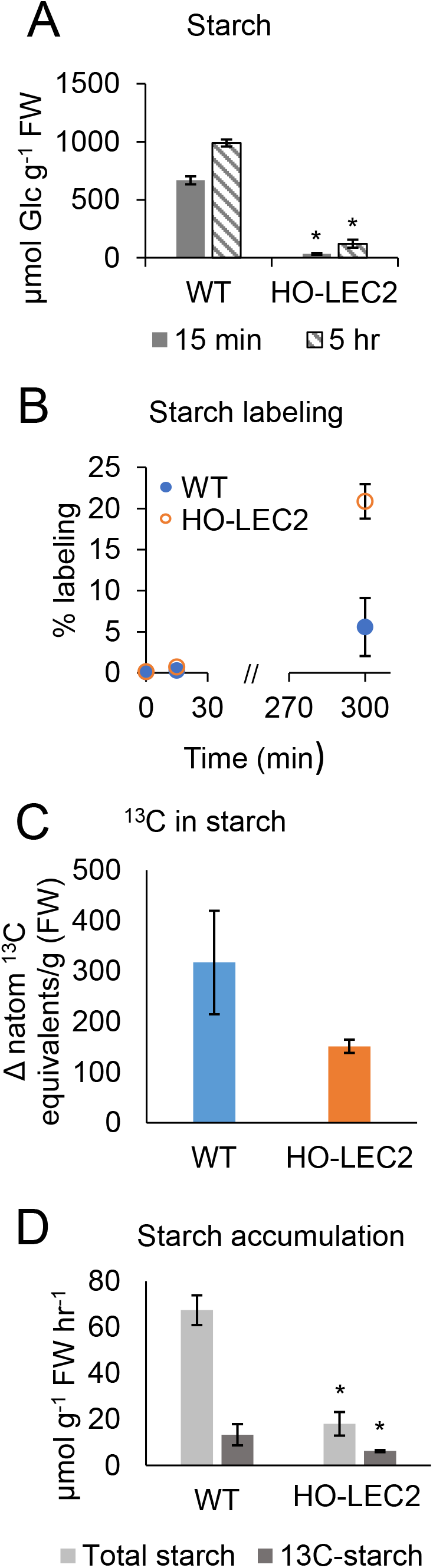
Starch production in WT and HO-LEC2 leaves over the 5 hr ^13^CO_2_-labeling experiment. The top expanded leaves of 62, 63, and 65 DAS WT and HO-LEC2 were provided with ^13^CO_2_ and sampled at 15 and 300 min timepoints. A, Total starch levels at 15 and 300 min. Asterisks indicate significant differences from WT in the HO-LEC2 line (Student’s *t*-test, *P* < 0.05). B, Average % ^13^C label incorporation (corrected for natural abundance). C, Amount of ^13^C in starch produced over the labeling period. ^13^C amounts (n atom ^13^C equivalents g^-1^ FW) were calculated as previously described (Arrivault et al., 2016). D, Differences in total starch production rates (light gray bars) and ^13^C-labeled starch production rates (dark gray bars). ^13^C-labeled starch amounts were calculated by summing the labeled isotopologue amounts determined by multiplying isotopologue abundance with the average pool size of starch-digested glucose. The results given are mean ± SD of three biological replicates. Asterisks indicate significant differences from WT in the HO-LEC2 line (Student’s *t*-test, *P* < 0.05).

Though the reduced starch production was consistent with the postulated tradeoff of carbon for lipid biosynthesis, the interpretation of labeling data for other metabolites was more complicated. Fluxes of labeled carbon through central carbon metabolites were estimated by calculating the amount of ^13^C in the metabolites (Fig.7) as previously described (Arrivault et al., 2016), in addition to considering average ^13^C-enrichment (Supplemental Fig.S3) and metabolite pool size data (Supplemental Fig.S4) (Szecowka et al., 2013; Ma et al., 2014). Calculation of n atom ^13^C equivalents g^-1^ FW enables labeling comparisons between WT and HO-LEC2 metabolites while bypassing potentially confounding differences in metabolite pool sizes and contribution of inactive pools between WT and HO-LEC2. Isotopic labeling resulted in some expected differences in fluxes through metabolic pathways between WT and HO-LEC2 leaves. For example, estimated ^13^C fluxes through the CBC/pentose phosphate pathway intermediates sedoheptulose-7-phosphate and dihydroxyacetone phosphate were lower in HO-LEC2 leaves (Fig.7; Supplemental Fig. S3), consistent with reduced carbon assimilation and decreased total vegetative biomass observed in the greenhouse. However, the movement of labeled carbon through hexose phosphates including the starch glucosyl donor ADP-glucose, the sucrose precursor UDP-glucose, and glucose 6-phosphate was higher in the HO-LEC2 line compared to WT (Fig.7; Supplemental Fig. S3). ADP-glucose reached a near-maximal ^13^C-enrichment of ~70% after only 15 min of labeling in both WT and HO-LEC2 leaves (Supplemental Fig.S3), indicating the majority of the ADP-glucose pool is labeled by an early time and potentially suggesting that labeled starch production over the observed experimental time should be similar between lines. The lack of further ^13^C increase in ADP-glucose after 15 min of ^13^CO_2_-labeling in WT leaves combined with the higher amount of the unlabeled isotopologue of ADP-glucose remaining in WT compared to HO-LEC2 after five hours of labeling (Supplemental Fig.S5A-B), however, is consistent with the increased production of unlabeled starch over the labeling period observed in WT. Levels of sucrose, UDP-glucose, and ADP-glucose did not change significantly over the course of the labeling experiment for either WT or HO-LEC2 (Supplemental Fig.S4), suggesting that no major changes in partitioning towards sucrose or starch occurred over the course of the labeling period. The combined data thus suggest a more complicated spatial separation of labeled and unlabeled metabolite intermediates.

**Figure 7.**
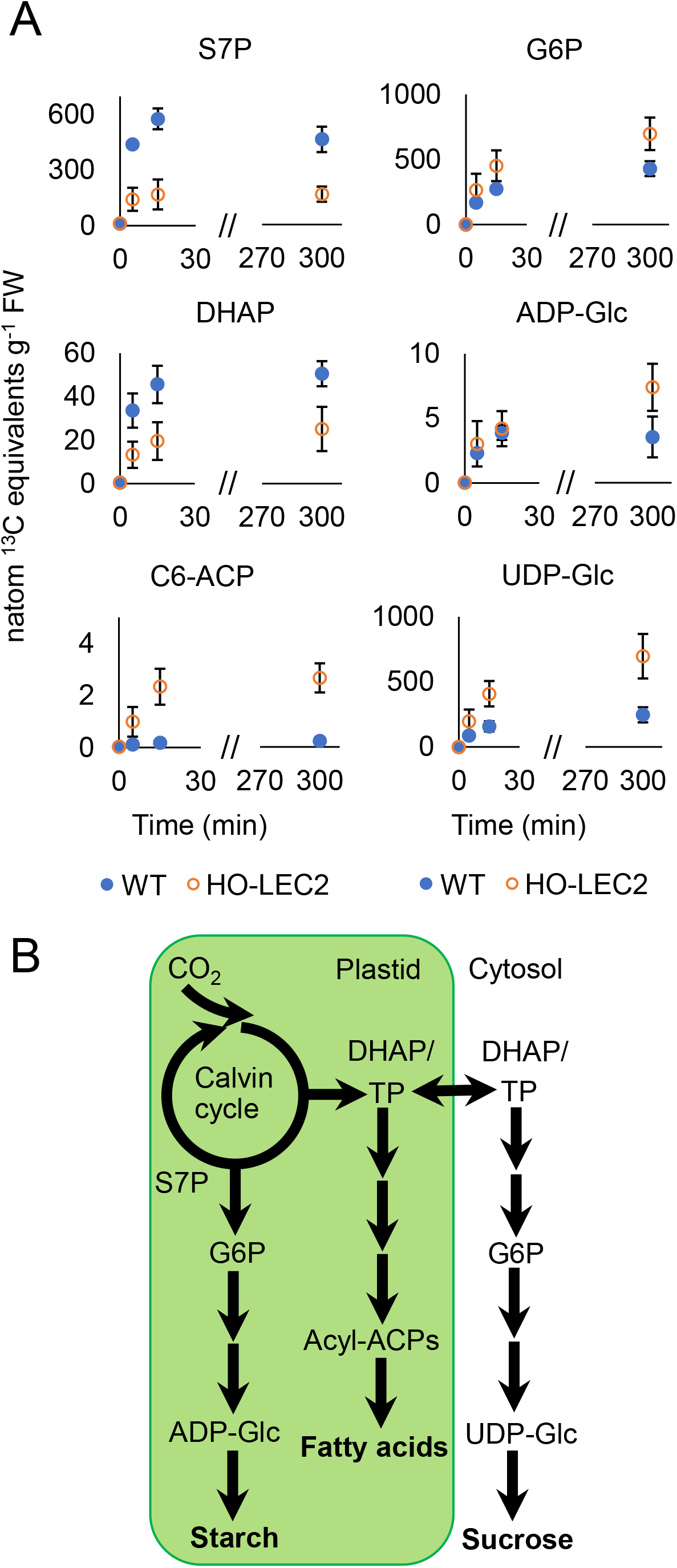
Visualization of carbon flux in high-oil accumulating leaves. A, Labeling kinetics of sedoheptulose 7-phosphate (S7P), dihydroxyacetone phosphate (DHAP), C6-ACP, glucose 6-phosphate (G6P), ADP-glucose (ADP-Glc), and UDP-glucose (UDP-Glc) presented as amount of ^13^C present in these metabolites over the labeling period. WT labeling patterns are marked in blue and HO-LEC2 patterns in orange. The results given are mean ± SD of three biological replicates. B, Metabolic map with outputs of interest in bold. Two arrows in a single step indicate two or more enzymatic reactions.

### Isotopically labeled acyl-ACPs indicate cellular heterogeneity in tobacco leaves

The ^13^CO_2_ transient labeling experiment also provided a unique opportunity to inspect the fatty acid biosynthesis process through the detection and quantification of isotope incorporation into acyl-ACPs. The multiple reaction monitoring LC-MS/MS method was adjusted to include transitions for all isotopologues of the different length acyl-ACPs, with C6-ACP reported as representative. Dynamic movement of labeled carbons through the acyl portions of acyl-ACPs were higher in HO-LEC2 leaves compared to WT (Fig.7; Supplemental Fig.S3), indicating increased flux through fatty acid synthesis. These findings thus provide dynamic evidence for elevated lipid biosynthetic rates in 63 DAS HO-LEC2 leaves that is consistent with FAME data. Labeling in WT acyl-ACPs leveled off at an average ^13^C-enrichment of 20% by five hours (Supplemental Fig.S3), indicating a significant portion of carbon flux to acyl-ACPs does not come directly from the CBC in WT plants or that a pool of acyl-ACPs remains unlabeled and is not a part of ‘active’ metabolism (Allen and Young, 2020). Analysis of mass isotopologue distribution changes of the acyl chains suggests the involvement of two differentially labeled acetyl-CoA pools since changes in the proportions of the different labeled isotopologues over time do not indicate linear incorporation of label (Supplemental Fig.S5C-D). One possibility is that WT leaves are deriving a greater portion of their acetyl-CoA from the large pool of relatively unlabeled cytosolic or vacuolar citrate (Rangasamy and Ratledge, 2000) in addition to plastidic pyruvate compared to HO-LEC2. The involvement of other subcellular pools such as mitochondrial acyl-ACPs, though thought to be small, also cannot be completely ruled out (Chuman and Brody, 1989; Gueguen et al., 2000). The increased stepwise labeling patterns of HO-LEC2 acyl-ACPs indicate that ^13^C label is incorporated more directly into fatty acid synthesis, likely due to elevated expression of chloroplastic isoforms of lower glycolytic enzymes (Vanhercke et al., 2017). The findings from multiple isotopic measurements of incomplete labeling of metabolites such as ADP-glucose and acyl-ACPs thus suggest cellular heterogeneity in tobacco leaves, with the effects of spatial segregation more pronounced in the WT line.

## Discussion

There have been many efforts to increase lipid levels in the vegetative tissues of plants (reviewed in Vanhercke et al., 2019b), with some of the most impressive increases in TAG occurring in tobacco (Vanhercke et al., 2014b; Vanhercke et al., 2017). TAG yields in transgenic tobacco leaves have far surpassed levels achieved in other plant species, even when similar transgene combinations are manipulated. In the current study, leaves from WT and the high oil-producing HO-LEC2 transgenic tobacco line were metabolically characterized to explore why tobacco is so amenable to foliar lipid engineering. The consequences of lipid production on dynamic photosynthetic carbon partitioning were surveyed in plants across development to examine the age when the synthesis rather than the accumulation of lipids was greatest. We measured lipid levels of WT and HO-LEC2 leaves at several plant ages before the onset of flowering and observed elevated foliar lipid levels at all ages (Fig.1B), including young HO-LEC2 plants a few days after transplantation. Further dramatic increases in TAG accumulation by nine weeks after planting likely reflect the construct design with overexpression of the *LEC2* gene under the control of the *A. thaliana* SAG12 senescence-specific promoter (Vanhercke et al., 2017). We targeted leaves from plants at the nine-week plant age in subsequent analyses due to the rapid change in lipids at this time.

### Isotopically labeled acyl-ACPs provide a means to assess the dynamics of fatty acid biosynthesis in the high oil line

An often overlooked aspect in the assessment of metabolism is that accumulation over time reflects both biosynthesis and metabolic turnover events (Allen, 2016a; Allen, 2016b). Frequently the accumulated level of a storage reserve such as lipid can be measured easily, but such measures do not capture the amount that may be turned over and can underestimate biosynthesis if turnover is not negligible. Acyl-ACP pools are specific components to the fatty acid biosynthetic process and represent a conclusive measure of fatty acid production. We paired isotopic labeling with a recently developed acyl-ACP quantification method to assess the dynamics of fatty acid metabolism. Our results provide evidence of elevated fluxes through fatty acid synthesis in HO-LEC2 leaves at the nine-week stage that validate the observed increase in lipid accumulation (Fig.2; Fig.7). The elevated daytime pools of HO-LEC2 acyl-ACPs indicate that most of the increased lipid production occurs during the day like starch, consistent with the increased activity in the light of the acetyl-CoA carboxylase enzyme responsible for the first committed step in fatty acid synthesis (Browse et al., 1981; Sauer and Heise, 1984; Sasaki et al., 1997). The lower lipid production in WT plants uncovered incomplete acyl-chain labeling patterns that suggest increased metabolic heterogeneity in WT leaves, a hypothesis supported by labeling data from other metabolites.

### Unanticipated differences in estimated carbon fluxes through ADP-glucose and starch reflect spatiotemporal differences in metabolism

No decreases in ^13^C content, ^13^C-enrichment, or pool sizes of ADP-glucose, UDP-glucose, or their common glucose 6-phosphate precursor were observed in HO-LEC2 leaves compared to WT at the nine-week age (Fig.7; Supplemental Fig.S3, Supplemental Fig.S4). The labeling results suggest that rates of HO-LEC2 photosynthetic carbohydrate biosynthesis during the day are higher than or similar to WT despite the increased lipid synthesis in HO-LEC2 at this stage. However, measurements of total and ^13^C-labeled starch accumulation as well as ^13^C content in starch over a five-hour labeling period reveal lower rates of starch production in HO-LEC2 leaves (Fig.6). In addition, the large differences between total and labeled starch production suggest the synthesis of a substantial amount of unlabeled starch in distinct intercellular locations during the labeling period (Fig.8). The unchanging ^13^C content in ADP-glucose after 15 min of ^13^CO_2_-labeling in WT leaves (Fig.7) and the significant proportion of unlabeled ADP-glucose remaining in WT leaves after five hours of labeling (Supplemental Fig.S5A) are consistent with this hypothesis, along with a number of observations in the literature. For example, tobacco bundle-sheath cells can utilize carbon from the vasculature rather than the stomata to produce starch (Hibberd and Quick, 2002) that can be significant relative to production in the mesophyll cells for some C3 plants (Miyake and Maeda, 1976; Miyake, 2016), possibly accounting for the unlabeled fraction (Fig.8C). Turnover of glandular trichome metabolites including acyl-sugars and diterpenes that represent 10-15% of leaf dry weight in certain *Nicotiana* species (Fobes et al., 1985; Wagner, 1991; Wagner et al., 2004; Slocombe et al., 2008) can also provide a source of unlabeled carbon for both lipid and carbohydrate production (Fig.8B), reflected by the unlabeled sugar phosphate and acyl-ACP fractions in leaves. Furthermore, Rubisco in glandular trichome cells is specialized to refix CO_2_ released during the synthesis of secondary metabolites (Laterre et al., 2017) and can provide additional pools of unlabeled primary metabolites. The lower proportion of unlabeled metabolites in HO-LEC2 leaves relative to WT (Supplemental Fig.S5) suggests that the increased push of carbon flux into TAG synthesis results in reduced metabolic pathway heterogeneity, with diminished production and turnover of secondary metabolites a likely consequence.

**Figure 8.**
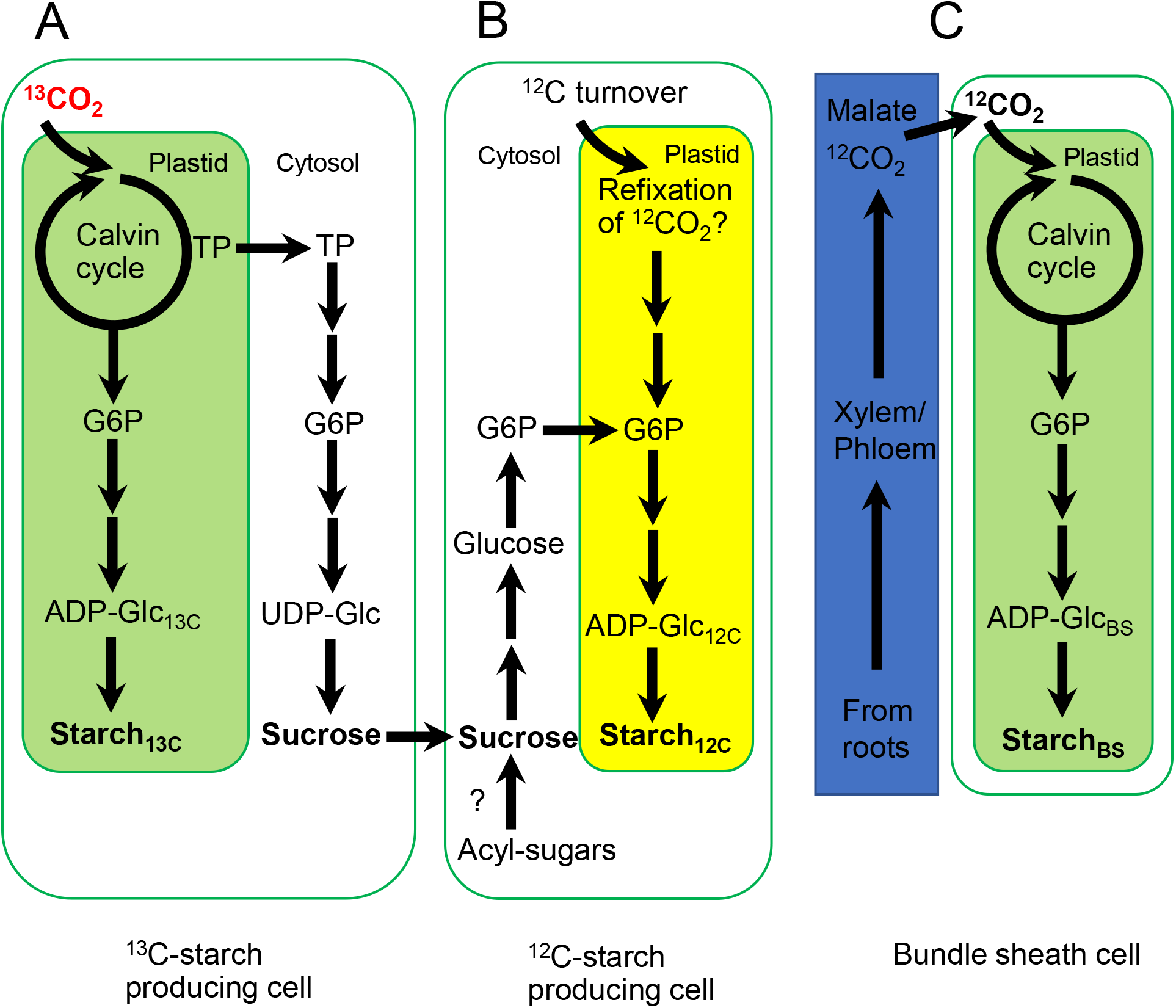
Putative map of carbon flux through heterogenous cells that produce both labeled and unlabeled starch within a photosynthetic leaf. ^13^C-labeling of starch pools in A, photosynthetic source cells would be diluted by B and C, production of unlabeled starch in cells relying on exported sucrose and/or able to use other unlabeled sources of carbon.

### Carbon from non-transient starch is redirected to lipid synthesis in high oil accumulating leaves

At the nine-week age, HO-LEC2 leaves display a drop in total starch that coincides with enhanced lipid production, suggesting a tradeoff in carbon. Our results demonstrate that WT tobacco leaves accumulate large amounts of non-transient starch with plant age, thus providing a ready pool of stored carbon in the chloroplasts for lipid synthesis. This starch can account for the majority of the carbon from the increased lipids in HO-LEC2 leaves (Fig.5), with the remaining carbon difference possibly due to decreased levels of secondary metabolites. Non-transient starch accumulation is absent in older HO-LEC2 plants, with diurnal starch patterns resembling those of young WT leaves. The lack of an excess starch pool in young HO-LEC2 plants that are still producing large amounts of lipid explains the hindered growth during establishment. In the high oil parent tobacco line lacking overexpression of the HO-LEC2 gene (Vanhercke et al., 2014b), diurnal starch levels and patterns in vegetative leaves are similar to WT (Mitchell et al., 2020), and the plants also accordingly suffer significant growth penalties. The redirection of carbon from starch to lipid has been previously demonstrated in Arabidopsis leaves via the combination of lipid engineering with mutants deficient in starch synthesis (Sanjaya et al., 2011; Zhai et al., 2017; Yu et al., 2018), though lipid levels did not come close to those achieved in transgenic tobacco leaves. Thus, the large non-transitory starch pools present in older tobacco leaves provide a possible explanation as to why leaf lipid engineering efforts have proven most successful in tobacco. Since Arabidopsis starch levels are normally tightly regulated for carbon use efficiency, it is expected that relatively little carbon can be shunted from starch to lipid without drastically affecting plant growth and viability, possibly explaining the more modest increases in TAG yields obtained in transgenic Arabidopsis plants with the same genes targeted as the high oil tobacco lines (Kelly et al., 2013; Zhai et al., 2017). A recent study engineering similar gene combinations in sorghum leaves (Vanhercke et al., 2019a) has also fallen short of the high lipid yields achieved in tobacco, further emphasizing the importance of a pre-existing carbon pool in achieving high foliar TAG yields.

In addition to carbon, the repartitioning of starch could provide reducing equivalents needed for fatty acid biosynthesis in HO-LEC2 plants. The utilization of glucose from starch breakdown might involve hexose phosphate processing through the oxidative pentose phosphate (OPP) pathway to produce NADPH, CO_2_, and ribose phosphates. Though operation of the CBC and the OPP pathway in tandem create a futile cycle, limited OPP activity later in development when *LEC2* is active could utilize stored starch to replenish CBC intermediates and reducing equivalents consumed in fatty acid synthesis. There is evidence that operation of the oxidative and reductive steps in pentose phosphate pathways are not mutually exclusive and that some operation of both may be important in particular circumstances (Sharkey and Weise, 2016; Li et al., 2019; Preiser et al., 2019; Weise et al., 2019; Allen and Young, 2020).

### What is the role of non-transient starch in WT tobacco?

A question raised by our findings concerns the unknown role of the non-transient foliar starch pool in WT plants. Since high oil-producing HO-LEC2 plants are capable of producing large amounts of viable seed, remobilization of this starch does not appear to be necessary for normal development. As the production of a significant, apparently unused storage compound is energetically inefficient and sequesters carbon that could otherwise be utilized for growth, defense, or reproduction, it is likely that that foliar starch plays a role besides that of a nighttime carbon source in older tobacco leaves. One hypothesis is that this starch pool acts as an important buffer against various environment stresses such as prolonged cloudy days by providing a readily accessible reserve carbon source. Starch levels have been shown to change in response to abiotic stresses (Weise et al., 2006; Thalmann et al., 2016; Thalmann and Santelia, 2017), with starch levels decreasing in response to most stresses but increasing during high salinity and cold temperature. The enzymes mediating starch degradation induced by stress were also often found to be different than the enzymes required for night-time starch breakdown, indicating a complex degree of regulatory control over starch metabolism. Further studies probing the function(s) of non-transient foliar starch are warranted, considering that starch metabolism does not appear to be consistent across different plant species and organs (Smith and Zeeman, 2020).

## Concluding Remarks

We have shown tobacco leaves from older WT plants are metabolically acting as dual source-sink tissues, assimilating carbon for local cellular requirements, synthesizing sucrose for export, and producing and storing non-transient starch. Knocking down starch biosynthetic genes such as AGPase in high starch-accumulating plants like tobacco may have presented an obvious next target for further optimization of foliar lipid production, but HO-LEC2 leaves are already redirecting significant carbon from starch to lipid. Since younger HO-LEC2 plants showed delayed growth and lower lipid accumulation when foliar starch was more transient, the amount of lipid that can be produced was limited by the energetic costs required for daily plant life. The continued observation of diurnal starch patterns across development in both WT and HO-LEC2 leaves suggests some basal level of transient starch is necessary for proper plant growth and maintenance. The developmental accumulation of non-transient starch in older tobacco plants allows for increased metabolic plasticity of carbon sources, with HO-LEC2 leaves simply substituting one form of stored carbon (starch) for another (TAGs). Plant species like *Arabidopsis thaliana* that finely regulate starch metabolism such that most starch reserves are depleted at dawn do not maintain a pool of “excess” available plastidic carbon for lipid production in leaves, possibly explaining the more limited lipid yield increases obtained in other plant species when the same gene combinations are manipulated (Vanhercke et al., 2019b). Thus, future efforts to engineer lipid production in vegetative tissues will need to consider the current capacity of the crop to store starch or other carbon reserves in leaves or stems. Sugarcane offers an encouraging case due to its accumulation of sucrose within internode stalks, but efforts thus far to engineer increased lipid accumulation have not yet matched the yields achieved in tobacco leaves (Zale et al., 2016; Parajuli et al., 2020), despite the relatively larger sink potential of stem tissue compared to leaves. It is likely that localization of the tobacco starch carbon pool in chloroplasts contributes to a more direct switch to oil production than the sugarcane sucrose pool compartmentalized in vacuoles and apoplast. Our findings indicate that a better understanding of plants that accumulate large non-essential “sink” pools of carbon-rich metabolites in their vegetative tissues could provide promising new platforms for vegetative oil crops.

## Materials and Methods

### Plant growth and leaf collection

WT (Wisconsin 38) and HO-LEC2 transgenic *Nicotiana tabacum* plants described previously (Vanhercke et al., 2017) were germinated in a greenhouse at 27°C/25°C (day/night) with 60-90% humidity before being transplanted into larger pots at three weeks after sowing. Plants were grown under 12 h/12 h light/dark conditions, with supplemental lighting activating whenever light readings dropped below 500 μmol*m^-2^s^-1^. Multiple sets of leaf punches (8.5 mm in diameter) were collected from the apexes of the top expanded leaf of four plants of each genotype at 26, 41, 55, 62, and 72 days after sowing (DAS). Sample collection was at noon unless otherwise indicated. The top expanded leaf for WT plants was leaf 8 at 41 DAS, leaf 11 at 55 DAS, leaf 13 at 63 DAS, and leaf 15 at 72 DAS. The top flat leaf, though not yet fully expanded, was leaf 6 for WT plants at 26 DAS. The top expanded leaf for HO-LEC2 plants was always one leaf below that of WT due to delayed initial growth. The collected leaf discs were frozen in liquid nitrogen and stored at −80°C until further processing. For starch and lipid assays, leaf discs were further lyophilized for 2 days at −50°C on a Labconco FreeZone freeze dryer before extraction.

### Lipid measurements

Total lipid levels were measured using a protocol modified from Allen and Young, 2013. Briefly, C15:0 and C17:0 triacylglycerol internal standards were added to 10-15 mg of lyophilized leaf tissue along with freshly prepared 5% sulfuric acid:methanol (v/v) and 0.2% butylated hydroxytoluene (BHT) in methanol. Samples were vortexed and heated at 110°C for 3 hrs, vortexing hourly. After cooling to room temperature, 0.9% NaCl (w/v) was added to each sample to quench the reaction and fatty acid methyl esters (FAMEs) were extracted using hexanes. FAMEs were quantified by gas chromatography-flame ionization (GC-FID) using a DB23 column (30 m, 0.25-mm i.d., 0.25-μm film; J&W Scientific), with the GC operated in split mode (15:1). The FID was run with an oven program that started at 180°C for 1 min, ramped from 180°C to 260°C at 20°C per min, held at 260°C for 7 mins, and ended with a half minute equilibration. Comparisons of peak areas to the two internal standards were used for quantification. Carbon compositions of 77.8% and 77.2% were determined for WT and HO-LEC2 TAGs respectively, utilizing their respective measured fatty acid distributions and formulas, when calculating biomass as carbon.

### Starch assay

The total concentration of foliar starch was determined by using the Megazyme starch assay kit (Megazyme International Ireland), using the AOAC Official Method 996.11 (Approved Methods of the AACC citation) modified to adjust the final assay volume for reading on a 96-well plate reader. Briefly, 10-15 mg of lyophilized leaf tissue was powdered and washed twice with 80% ethanol at 85°C before being heated with DMSO for 10 min at 110°C. The samples were then treated with α-amylase at 110°C for 12 minutes (vortexing every 4 min) followed by amyloglucosidase at 50°C for 1 hour. After centrifugation, serial dilutions of the supernatant were incubated with the GOPOD reagent at 50°C for 20 min before absorbance at 510 nm was measured against reagent blanks. Sample concentrations were determined using a standard curve generated with maize starch standards processed alongside leaf samples, with an additional standard curve generated with glucose standards used to check assay recovery. An additional acyl-sugar extraction in 1 mL isopropanol:acetonitrile:water (3:3:2, v/v/v) containing 0.1% formic acid by volume with gentle rocking for 5 min (Ning et al., 2015) inserted after the 80% ethanol washes on a subset of samples did not change final starch measurements compared to normally extracted samples from the same leaves. A carbon composition of 42.9% was used for starch when calculating biomass as carbon, utilizing the chemical formula for maltotriose.

### Metabolite extraction and quantification

Soluble sugars and sugar phosphates were extracted using a protocol modified from Ma et al., 2017, taking care to keep samples chilled throughout. 1 mL chilled methanol:chloroform (7:3) and 20 μL of 100 μg/mL PIPES/norvaline/ribitol internal standard mix were add to 100 mg (FW) leaf samples, which were then powdered using a ball mill at 29 Hz/sec for 5 mins and incubated on a mixer at 4°C for 2 hrs. Aqueous metabolites were extracted with chilled water before being centrifuged to separate phases. The upper aqueous phase was collected and dried in a LabConco CentriVap at 25°C before resuspending in 100 μL 50% methanol. The samples were passed through a 0.45 μm spin filter (Costar^®^, Corning Inc.) before further analysis on LC-MS/MS. Samples were injected on an Infinity Lab Poroshell 120 Z-HILIC column (2.7 μm, 100 x 2.1 mm; Agilent technologies, Santa Clara, CA, USA). All LC-MS/MS analyses were performed using a Shimadzu (UFLC_XR_) HPLC system connected to a tandem triple quadrupole mass spectrometer (6500 QTRAP^®^ LC-MS/MS system, Applied Biosystems) equipped with a Turbo V^™^ electrospray ionization (ESI) source. Analyst software (version 1.6.2) (AB SCIEX, Concord, Canada) was used to collect and analyze data. Ions were detected and monitored using a targeted multiple-reaction monitoring (MRM) approach with the parameters as previously described (Czajka et al., 2020).

Sugar phosphates were eluted and run using parameters as previously described (Czajka et al., 2020). Soluble sugars were eluted using acetonitrile: 20 mM ammonium formate (95:5 v/v) (A) and 20 mM ammonium formate in water (B), with both buffers adjusted to pH 6.8. A flow rate of 0.2 mL/min was used with a gradient of 95 to 60% B over 3.5 minutes, then to 25% B over the next 2 min, a hold at 25% B for 1.5 minutes, a return to 95% B over a minute, and a final re-equilibration for 6 minutes. Sugars were run in negative ion mode, with the curtain gas set to 35 psi, the ion spray voltage to 4.5 kV, the source temperature to 400°C, the nebulizing gas (GS1) to 40 psi, the focusing gas (GS2) to 40 psi, and the entrance potential to 10 V. The value for entrance potential was default (−10) for all analytes. Other parameters for MRMs were optimized using direct injections of individual sugar standards. Metabolite concentrations and recoveries were calculated based on external calibration curves and internal standards (PIPES for sugar phosphates and ribitol for sugars). A 1/*x*^2^ weighting factor was applied for external calibration curves as suggested by Gu et al., 2014.

Protein quantification was performed by directly hydrolyzing the protein from leaf samples and quantifying the resulting individual amino acids as previously described (Kambhampati et al., 2019). Protein hydrolysis was performed in 4 M methanesulfonic acid with 0.2% tryptamine and 20 uL of 1 mM ^13^C^15^N labeled amino acid standard mix for 22 hrs at 110°C. Samples were neutralized with 4 M sodium hydroxide before being dried in the CentriVap, resuspended in 50% methanol, and passed through 0.8 μM PES spin filters (Vivaclear, Sartorius Stedim Biotech) before being run on the LC-MS/MS. Amino acids were eluted using the same HILIC column as above using acetonitrile: 20 mM ammonium formate (90:10 v/v) (A) and 20 mM ammonium formate in water (B), with both buffers adjusted to pH 3.0. A flow rate of 0.25 mL/min was used with a gradient of 95 to 60% B over 4.5 minutes, 60 to 25% B over the next 1.5 min, a hold at 25% B for 1 min, a return to 95% B over 1.75 min, and a final re-equilibration for 5.25 min. Amino acids were run in positive ion mode, with the curtain gas set to 35 psi, the ion spray voltage to 4.5 kV, the source temperature to 400°C, the nebulizing gas (GS1) to 40 psi, the focusing gas (GS2) to 30 psi, and the entrance potential to 10 V. Other parameters for MRMs were optimized using direct injections of individual amino acid standards. Amino acid concentrations and recoveries were calculated based on direct comparison of peak areas with their respective labeled internal standards as previously described (Kambhampati et al., 2019).

### ^13^CO_2_ labeling

^13^CO_2_ labeling studies were performed on greenhouse-grown WT and HO-LEC2 plants. The tips of the top expanded leaves of 62-64 DAS WT and HO-LEC2 plants (leaves 13 and 12, respectively) were clamped in custom-made transparent cylindrical labeling chambers (vol.= 50.6676 cm^3^) for labeling using premixed gas containing ^13^CO_2_ at a ^13^CO_2_/N2/O2 ratio of 0.033:78:21.967 (Aldrich) with a flow rate of 0.5 L/min. The leaf was positioned in the labeling chamber such that roughly even amounts of tissue were present on either side of the midvein with a small gap left on the side of the leaf closest to the exit hole opposite the gas inlet. The portion of the leaf within the chamber was marked during labeling. After providing leaves with label for either 5, 15, or 300 minutes, the leaf was removed from the labeling chamber and rapidly quenched using a freeze-clamp prechilled in liquid nitrogen. The frozen leaf sections were then dropped onto a prechilled metal plate where the labeled leaf section was isolated before being coarsely ground and stored at −80° C. The same procedures were carried out in parallel on the top expanded leaves of unlabeled control plants. All labeling experiments were carried out from late morning to mid-afternoon (10 AM-4 PM), with a WT and a HO-LEC2 plant being labeled simultaneously for a total of three plants of each genotype being labeled on successive days. After extracting sugars and sugar phosphates, their mass isotopologue distributions were determined via LC-MS/MS (Ma et al., 2017), with correction for natural ^13^C abundance performed as previously described (Fernandez et al., 1996).

Labeled starch isotopologues were calculated by multiplying isotopologue abundance with the average pool sizes of starch-digested glucose measured with the Megazyme kit. ^13^C starch content was determined by summing the amounts of labeled isotopologues as previously described (Arrivault et al., 2016; Fernandez et al., 2017), except the analysis of labeled isotopologues was directly performed on glucose from α-amylase- and amyloglucosidase-digested starch. The amount of ^13^C in metabolites was determined by multiplying the calculated amount of each labeled isotopologue by the number of labeled carbons in each isotopologue and summing the results as previously described (Arrivault et al., 2016).

### Acyl-ACP extraction

Acyl-ACPs were precipitated using trichloroacetic acid (TCA) using a protocol as previously described (Nam et al., 2020). In brief, 5% TCA was added to 30 or 60 mg of ground fresh leaf tissue for WT and HO-LEC2, respectively. After vortexing and pelleting the protein via centrifugation, the supernatant was removed and the pellet reprecipitated with 1% TCA. After resuspending the pellet in 50 mM MOPS (pH 7.5) and incubating on ice for 1 hr, cellular debris was removed via centrifugation. TCA was added to the supernatant to a final concentration of 10% and incubated at −80°C overnight before thawing on ice and pelleting via centrifugation. After washing the ACP protein pellet with 1% TCA and a final centrifugation step, the pellet was resuspended in minimal volume of 50 mM MOPS (pH 7.5) before being digested with Asp-N endoproteinase (Sigma-Aldrich) at 1:50 (enzyme:acyl-ACP) at 37°C for 2.5 hrs. The reaction was quenched by the addition of methanol to a final concentration of 50%, and the samples were passed through 0.45 um spin filters (Costar^®^, Corning Inc.) before analysis on LC-MS/MS.

The AspN-digested acyl-ACP products were separated on a reversed-phase column (Discovery^®^ BIO Wide Pore C18; Sigma-Aldrich, 10 cm, 2.1 mm, 3 mm) using acetonitrile:10 mM ammonium formate (90:10 v/v) (A) and acetonitrile:10 mM ammonium formate (90:10 v/v) (B), with both buffers at pH 3.5. A flow rate of 0.2 mL/min was used with a gradient program from 100% A to 10% B over seven minutes, then to 100% B over the next 8 min, a hold for 4 min, a return to 0% B over 2 min, and a final re-equilibration for 9 min. Acyl-ACP samples were run in positive ion mode, with the curtain gas set to 20 psi, the ion spray voltage to 5.5 kV, the source temperature to 400°C, the nebulizing gas (GS1) to 40 psi, the focusing gas (GS2) to 35 psi, and the entrance potential to 10 V. The digested acyl-ACP molecules were identified using MRM parameters as previously described (Nam et al., 2020), with the addition of ^13^C-labeled isotopologues.

### Confocal Microscopy

Tissue samples were taken from the top expanded leaves of 67 DAS WT and HO-LEC2 plants. Leaf samples were stained for 45 minutes in aqueous BODIPY 505/515 (1 μg/ml), fixed in 4% paraformaldehyde in PBS, rinsed three times in water, and mounted in 6% agar. 100 μm thick leaf cross-sections were generated (Vibratome 1500 Sectioning System) for imaging using a Leica SP8-X confocal microscope with a 63 X HC Plan Apochromat water immersion objective (1.2 NA) and the pinhole set to 1 airy unit. The following excitation and emission wavelengths were used: 405 nm excitation and 415-450 nm emission for cell wall auto fluorescence (cyan), 488 nm excitation and 500-550nm emission for BODIPY-labeled lipid droplets (green), and 649 nm excitation and 659-779 nm emission for chlorophyll autofluorescence (magenta). Optical slices were collected at 1 μm z-intervals to create 3D maximum intensity projections.

### Data analysis

All calculations from raw data were done in Microsoft Excel, including graphing and statistical analysis with the aid of the Analysis ToolPak and Daniel’s XL Toolbox (version 7.3.4, by Daniel Kraus, Würzburg, Germany (www.xltoolbox.net)) add-ins.

## Supplemental Data

**Supplemental Figure S1.** Differences in fatty acid species in the top expanded leaf of 62 DAS WT and HO-LEC2 plants sampled at dawn and dusk.

**Supplemental Figure S2.** Soluble sugar levels in the top expanded leaf of WT and HO-LEC2 tobacco plants at 41, 55, 62, and 72 DAS across photoperiod.

**Supplemental Figure S3.** Percent ^13^C label incorporation (corrected for natural abundance) in sedoheptulose 7-phosphate (S7P), dihydroxyacetone phosphate (DHAP), C6-ACP, glucose 6-phosphate (G6P), ADP-glucose (ADP-Glc), and UDP-glucose (UDP-Glc) over the labeling period.

**Supplemental Figure S4.** Pool sizes of sedoheptulose-7-phosphate, dihydroxyacetone phosphate, C6-ACP, glucose 6-phosphate, ADP-glucose, and UDP-glucose at the start and end of the ^13^CO_2_ labeling experiment.

**Supplemental Figure S5.** Changes in mass isotopologue distributions (MIDs) of ADP-Glc and C6-ACP over the ^13^CO_2_ labeling experiment.

## Acknowledgements

We thank Dr. Xue-Rong Zhou (Commonwealth Scientific and Industrial Research Organisation (CSIRO)) for providing WT and HO-LEC2 transgenic tobacco seeds; Dr. Howard Berg (Integrated Microscopy Facility/Advanced Bioimaging Laboratory, Donald Danforth Plant Science Center) for preliminary microscopy efforts, Dr. Brad Evans and Jon Mattingly (Proteomics and Mass Spectrometry Facility, Donald Danforth Plant Science Center) for their assistance with LC-MS instrumentation; and Kevin Reilly, Kris Haines, Matt Adams, and Aileen Wok (Integrated Plant Growth Facility, Donald Danforth Plant Science Center) for their assistance with plant growth and propagation. We also acknowledge imaging support from the Advanced Bioimaging Laboratory at the Danforth Plant Science Center. The Leica SPX-8 confocal microscope used was acquired through National Science Foundation Major Research Instrumentation grant (DBI-1337680) while the QTRAP LC-MS/MS used was acquired through (DBI-1427621). The authors acknowledge support for aspects of this work from United States Department of Agriculture-Agricultural Research Station, United States Department of Agriculture-National Institute of Food and Agriculture grant awards (2017-67013-26156 and 2016-67013-24585), National Science Foundation awards (DBI-1616820 and DBI-1828365), and National Institutes of Health award (U01 CA235508).

## Literature cited

Ainsworth EA, Rogers A, Leakey ADB, Heady LE, Gibon Y, Stitt M, Schurr U (2007) Does elevated atmospheric [CO2] alter diurnal C uptake and the balance of C and N metabolites in growing and fully expanded soybean leaves? J Exp Bot 58: 579–591

Allen DK (2016a) Assessing compartmentalized flux in lipid metabolism with isotopes. Biochim Biophys Acta - Mol Cell Biol Lipids 1861: 1226–1242

Allen DK (2016b) Quantifying plant phenotypes with isotopic labeling & metabolic flux analysis. Curr Opin Biotechnol 37: 45–52

Allen DK, Bates PD, Tjellström H (2015) Tracking the metabolic pulse of plant lipid production with isotopic labeling and flux analyses: Past, present and future. Prog Lipid Res 58: 97–120

Allen DK, Young JD (2020) Tracing metabolic flux through time and space with isotope labeling experiments. Curr Opin Biotechnol 64: 92–100

Allen DK, Young JD (2013) Carbon and nitrogen provisions alter the metabolic flux in developing soybean embryos. Plant Physiol 161: 1458–1475

Andrianov V, Borisjuk N, Pogrebnyak N, Brinker A, Dixon J, Spitsin S, Flynn J, Matyszczuk P, Andryszak K, Laurelli M, et al (2010) Tobacco as a production platform for biofuel: Overexpression of Arabidopsis DGAT and LEC2 genes increases accumulation and shifts the composition of lipids in green biomass. Plant Biotechnol J 8: 277–287

Arisz SA, Heo J-Y, Koevoets IT, Zhao T, van Egmond P, Meyer AJ, Zeng W, Niu X, Wang B, Mitchell-Olds T, et al (2018) DIACYLGLYCEROL ACYLTRANSFERASE1 Contributes to Freezing Tolerance. Plant Physiol 177: 1410 LP – 1424

Arrivault S, Obata T, Szecówka M, Mengin V, Guenther M, Hoehne M, Fernie AR, Stitt M (2016) Metabolite pools and carbon flow during C4 photosynthesis in maize: 13CO2 labeling kinetics and cell type fractionation. J Exp Bot 68: 283–298

Baerenfaller K, Massonnet C, Hennig L, Russenberger D, Sulpice R, Walsh S, Stitt M, Granier C, Gruissem W (2015) A long photoperiod relaxes energy management in Arabidopsis leaf six. Curr Plant Biol 2: 34–45

Bassham JA, Benson AA, Kay LD, Harris AZ, Wilson AT, Calvin M (1954) The Path of Carbon in Photosynthesis. XXI. The Cyclic Regeneration of Carbon Dioxide Acceptor. J Am Chem Soc 76: 1760–1770

Baud S, Lepiniec L (2010) Physiological and developmental regulation of seed oil production. Prog Lipid Res 49: 235–249

Beechey-Gradwell Z, Cooney L, Winichayakul S, Andrews M, Hea SY, Crowther T, Roberts N (2019) Storing carbon in leaf lipid sinks enhances perennial ryegrass carbon capture especially under high N and elevated CO2. J Exp Bot 71: 2351–2361

Bourgis F, Kilaru A, Cao X, Ngando-Ebongue G-F, Drira N, Ohlrogge JB, Arondel V (2011) Comparative transcriptome and metabolite analysis of oil palm and date palm mesocarp that differ dramatically in carbon partitioning. Proc Natl Acad Sci 108: 12527 LP – 12532

Browse J, Roughan PG, Slack CR (1981) Light control of fatty acid synthesis and diurnal fluctuations of fatty acid composition in leaves. Biochem J 196: 347–354

Cai Y, Whitehead P, Chappell J, Chapman KD (2019) Mouse lipogenic proteins promote the co-accumulation of triacylglycerols and sesquiterpenes in plant cells. Planta 250: 79–94

Caspar T, Huber SC, Somerville C (1985) Alterations in Growth, Photosynthesis, and Respiration in a Starchless Mutant of Arabidopsis thaliana (L.) Deficient in Chloroplast Phosphoglucomutase Activity. Plant Physiol 79: 11 –17

Caspar T, Lin TP, Kakefuda G, Benbow L, Preiss J, Somerville C (1991) Mutants of Arabidopsis with altered regulation of starch degradation. Plant Physiol 95: 1181–1188

Chuman L, Brody S (1989) Acyl carrier protein is present in the mitochondria of plants and eucaryotic micro-organisms. Eur J Biochem 184: 643–649

Czajka JJ, Kambhampati S, Tang YJ, Wang Y, Allen DK (2020) Application of Stable Isotope Tracing to Elucidate Metabolic Dynamics During Yarrowia lipolytica α-Ionone Fermentation. iScience 23: 100854

Diaz C, Lemaître T, Christ A, Azzopardi M, Kato Y, Sato F, Morot-Gaudry JF, Le Dily F, Masclaux-Daubresse C (2008) Nitrogen recycling and remobilization are differentially controlled by leaf senescence and development stage in Arabidopsis under low nitrogen nutrition. Plant Physiol 147: 1437–1449

Durand M, Mainson D, Porcheron B, Maurousset L, Lemoine R, Pourtau N (2018) Carbon source–sink relationship in Arabidopsis thaliana: the role of sucrose transporters. Planta 247: 587–611

Durrett TP, Benning C, Ohlrogge J (2008) Plant triacylglycerols as feedstocks for the production of biofuels. Plant J 54: 593–607

Fan J, Yan C, Roston R, Shanklin J, Xu C (2014) Arabidopsis Lipins, PDAT1 Acyltransferase, and SDP1 Triacylglycerol Lipase Synergistically Direct Fatty Acids toward β-Oxidation, Thereby Maintaining Membrane Lipid Homeostasis. Plant Cell 26: 4119 LP – 4134

Fan J, Yu L, Xu C (2017) A Central Role for Triacylglycerol in Membrane Lipid Breakdown, Fatty Acid β-Oxidation, and Plant Survival under Extended Darkness. Plant Physiol 174: 1517 LP – 1530

Fernandez CA, Des Rosiers C, Previs SF, David F, Brunengraber H (1996) Correction of 13C mass isotopomer distributions for natural stable isotope abundance. J Mass Spectrom 31: 255–262

Fernandez O, Ishihara H, George GM, Mengin V, Flis A, Sumner D, Arrivault S, Feil R, Lunn JE, Zeeman SC, et al (2017) Leaf starch turnover occurs in long days and in falling light at the end of the day. Plant Physiol 174: 2199–2212

Fobes JF, Mudd JB, Marsden MPF (1985) Epicuticular Lipid Accumulation on the Leaves of Lycopersicon pennellii (Corr.) D’Arcy and Lycopersicon esculentum Mill. Plant Physiol 77: 567–570

Geiger DR, Servaites JC (1994) Diurnal regulation of photosynthetic carbon metabolism in C3 plants. Annu Rev Plant Physiol Plant Mol Biol 45: 235–256

Geiger DR, Shieh Wen-Jang, Yu Xiao-Ming (1995) Photosynthetic carbon metabolism and translocation in wild-type and starch-deficient mutant Nicotiana sylvestris L. Plant Physiol 107: 507–514

Gibon Y, Pyl ET, Sulpice R, Lunn JE, HÖhne M, GÜnther M, Stitt M (2009) Adjustment of growth, starch turnover, protein content and central metabolism to a decrease of the carbon supply when Arabidopsis is grown in very short photoperiods. Plant, Cell Environ 32: 859–874

Graf A, Schlereth A, Stitt M, Smith AM (2010) Circadian control of carbohydrate availability for growth in Arabidopsis plants at night. Proc Natl Acad Sci U S A 107: 9458–9463

Graf A, Smith AM (2011) Starch and the clock: The dark side of plant productivity. Trends Plant Sci 16: 169–175

Gu H, Liu G, Wang J, Aubry AF, Arnold ME (2014) Selecting the correct weighting factors for linear and quadratic calibration curves with least-squares regression algorithm in bioanalytical LC-MS/MS assays and impacts of using incorrect weighting factors on curve stability, data quality, and assay perfo. Anal Chem 86: 8959–8966

Gueguen V, Macherel D, Jaquinod M, Douce R, Bourguignon J (2000) Fatty acid and lipoic acid biosynthesis in higher plant mitochondria. J Biol Chem 275: 5016–5025

Harrison CJ, Hedley CL, Wang TL (1998) Evidence that the rug3 locus of pea (Pisum sativum L.) encodes plastidial phosphoglucomutase confirms that the imported substrate for starch synthesis in pea amyloplasts is glucose-6-phosphate. Plant J 13: 753–762

Häusler RE, Schlieben NH, Schulz B, Flügge UI (1998) Compensation of decreased triose phosphate/phosphate translocator activity by accelerated starch turnover and glucose transport in transgenic tobacco. Planta 204: 366–376

Hibberd JM, Quick WP (2002) Characteristics of C4 photosynthesis in stems and petioles of C3 flowering plants. Nature 415: 451–454

Himelblau E, Amasino RM (2001) Nutrients mobilized from leaves of Arabidopsis thaliana during leaf senescence. J Plant Physiol 158: 1317–1323

Huber SC, Hanson KR (1992) Carbon partitioning and growth of a starchless mutant of Nicotiana sylvestris. Plant Physiol 99: 1449–1454

Ischebeck T, Krawczyk HE, Mullen RT, Dyer JM, Chapman KD (2020) Lipid droplets in plants and algae: Distribution, formation, turnover and function. Semin Cell Dev Biol. doi: https://doi.org/10.1016/j.semcdb.2020.02.014

Kambhampati S, Li J, Evans BS, Allen DK (2019) Accurate and efficient amino acid analysis for protein quantification using hydrophilic interaction chromatography coupled tandem mass spectrometry. Plant Methods 15: 46

Kaup MT, Froese CD, Thompson JE (2002) A role for diacylglycerol acyltransferase during leaf senescence. Plant Physiol 129: 1616–1626

Kelly AA, van Erp H V., Quettier AL, Shaw E, Menard G, Kurup S, Eastmond PJ (2013) The SUGAR-DEPENDENT1 lipase limits triacylglycerol accumulation in vegetative tissues of Arabidopsis. Plant Physiol 162: 1282–1289

Kichey T, Hirel B, Heumez E, Dubois F, Le Gouis J (2007) In winter wheat (Triticum aestivum L.), post-anthesis nitrogen uptake and remobilisation to the grain correlates with agronomic traits and nitrogen physiological markers. F Crop Res 102: 22–32

Koiwai A, Suzuki F, Matsuzaki T, Kawashima N (1983) The fatty acid composition of seeds and leaves of Nicotiana species. Phytochemistry 22: 1409–1412

Laterre R, Pottier M, Remacle C, Boutry M (2017) Photosynthetic trichomes contain a specific rubisco with a modified pH-dependent activity. Plant Physiol 173: 2110–2120

Li-Beisson Y, Shorrosh B, Beisson F, Andersson MX, Arondel V, Bates PD, Baud S, Bird D, DeBono A, Durrett TP, et al (2013) Acyl-Lipid Metabolism. Arab B 11: e0161

Li J, Weraduwage SM, Preiser AL, Tietz S, Weise SE, Strand DD, Froehlich JE, Kramer DM, Hu J, Sharkey TD (2019) A cytosolic bypass and g6p shunt in plants lacking peroxisomal hydroxypyruvate reductase1[open]. Plant Physiol 180: 783–792

Li W, Zhang H, Li X, Zhang F, Liu C, Du Y, Gao X, Zhang Z, Zhang X, Hou Z, et al (2017) Intergrative metabolomic and transcriptomic analyses unveil nutrient remobilization events in leaf senescence of tobacco. Sci Rep 7: 12126

Lin T-P, Caspar T, Somerville CR, Preiss J (1988) A Starch Deficient Mutant of Arabidopsis thaliana with Low ADPglucose Pyrophosphorylase Activity Lacks One of the Two Subunits of the Enzyme. Plant Physiol 88: 1175–1181

Lu Y, Gehan JP, Sharkey TD (2005) Daylength and circadian effects on starch degradation and maltose metabolism. Plant Physiol 138: 2280–2291

Ludewig F, Flügge UI (2013) Role of metabolite transporters in source-sink carbon allocation. Front Plant Sci 4: 231

Ma F, Jazmin LJ, Young JD, Allen DK (2014) Isotopically nonstationary 13C flux analysis of changes in Arabidopsis thaliana leaf metabolism due to high light acclimation. Proc Natl Acad Sci U S A 111: 16967–16972

Ma F, Jazmin LJ, Young JD, Allen DK (2017) Isotopically nonstationary metabolic flux analysis (INST-MFA) of photosynthesis and photorespiration in plants. *In* AR Fernie, H Bauwe, APM Weber, eds, Methods Mol. Biol. Springer New York, New York, NY, pp 167–194

Matheson N, Wheatley J (1962) Starch Changes in Developing and Senescing Tobacco Leaves. Aust J Biol Sci 15: 445

Matheson N, Wheatley J (1963) Diurnal-Nocturnal Changes in the Starch of Tobacco Leaves. Aust J Biol Sci 16: 70

Matt P, Schurr U, Klein D, Krapp A, Stitt M (1998) Growth of tobacco in short-day conditions leads to high starch, low sugars, altered diurnal changes in the Nia transcript and low nitrate reductase activity, and inhibition of amino acid synthesis. Planta 207: 27–41

McClain AM, Sharkey TD (2019) Triose phosphate utilization and beyond: From photosynthesis to end product synthesis. J Exp Bot 70: 1755–1766

Mengin V, Pyl ET, Moraes TA, Sulpice R, Krohn N, Encke B, Stitt M (2017) Photosynthate partitioning to starch in arabidopsis thaliana is insensitive to light intensity but sensitive to photoperiod due to a restriction on growth in the light in short photoperiods. Plant Cell Environ 40: 2608–2627

Mitchell MC, Pritchard J, Okada S, Zhang J, Venables I, Vanhercke T, Ral J (2020) Increasing growth and yield by altering carbon metabolism in a transgenic leaf oil crop. Plant Biotechnol J n/a: pbi.13363

Miyake H (2016) Starch Accumulation in the Bundle Sheaths of C3 Plants: A Possible Pre-Condition for C4 Photosynthesis. Plant Cell Physiol 57: 890–896

Miyake H, Maeda E (1976) Development of bundle sheath chloroplasts in rice seedlings. Can J Bot 54: 556–565

Nam J-W, Jenkins LM, Li J, Evans B, Jaworski JG, Allen DK (2020) A General Method for Quantification and Discovery of Acyl Groups Attached to Acyl Carrier Proteins in Fatty Acid Metabolism using LC-MS/MS. Plant Cell tpc.00954.2019

Niittylä T, Messerli G, Trevisan M, Chen J, Smith AM, Zeeman SC (2004) A Previously Unknown Maltose Transporter Essential for Starch Degradation in Leaves. Science (80-) 303: 87 LP – 89

Ölçer H, Lloyd JC, Raines CA (2001) Photosynthetic capacity is differentially affected by reductions in sedoheptulose-1,7-bisphosphatase activity during leaf development in transgenic tobacco plants. Plant Physiol 125: 982–989

Parajuli S, Kannan B, Karan R, Sanahuja G, Liu H, Garcia-Ruiz E, Kumar D, Singh V, Zhao H, Long S, et al (2020) Towards oilcane: Engineering hyperaccumulation of triacylglycerol into sugarcane stems. GCB Bioenergy. doi: 10.1111/gcbb.12684

Preiser AL, Fisher N, Banerjee A, Sharkey TD (2019) Plastidic glucose-6-phosphate dehydrogenases are regulated to maintain activity in the light. Biochem J 476: 1539–1551

Raines CA, Paul MJ (2006) Products of leaf primary carbon metabolism modulate the developmental programme determining plant morphology. J Exp Bot 57: 1857–1862

Rangasamy D, Ratledge C (2000) Compartmentation of ATP:citrate lyase in plants. Plant Physiol 122: 1225–1230

Sanjaya, Durrett TP, Weise SE, Benning C (2011) Increasing the energy density of vegetative tissues by diverting carbon from starch to oil biosynthesis in transgenic Arabidopsis. Plant Biotechnol J 9: 874–883

Sasaki Y, Kozaki A, Hatano M (1997) Link between light and fatty acid synthesis: thioredoxin-linked reductive activation of plastidic acetyl-CoA carboxylase. Proc Natl Acad Sci U S A 94: 11096–11101

Sauer A, Heise K-P (1984) Regulation of Acetyl-Coenzyme A Carboxylase and Acetyl-Coenzyme A Synthetase in Spinach Chloroplasts. Zeitschrift für Naturforsch C 39: 268–275

Severson R, Johnson A, Jackson D (1985) Cuticular constituents of tobacco: factors affecting their production and their role in insect and disease resistance and smoke quality. Rec Adv Tob Sci 11: 105–174

Sharkey TD, Weise SE (2016) The glucose 6-phosphate shunt around the Calvin-Benson cycle. J Exp Bot 67: 4067–4077

Slocombe SP, Cornah J, Pinfield-Wells H, Soady K, Zhang Q, Gilday A, Dyer JM, Graham IA (2009) Oil accumulation in leaves directed by modification of fatty acid breakdown and lipid synthesis pathways. Plant Biotechnol J 7: 694–703

Slocombe SP, Schauvinhold I, McQuinn RP, Besser K, Welsby NA, Harper A, Aziz N, Li Y, Larson TR, Giovannoni J, et al (2008) Transcriptomic and reverse genetic analyses of branched-chain fatty acid and acyl sugar production in Solanum pennellii and Nicotiana benthamiana. Plant Physiol 148: 1830–1846

Smith AM, Stitt M (2007) Coordination of carbon supply and plant growth. Plant, Cell Environ 30: 1126–1149

Smith AM, Zeeman SC (2020) Starch: A Flexible, Adaptable Carbon Store Coupled to Plant Growth. Annu Rev Plant Biol 1–29

Stitt M, Quick WP (1989) Photosynthetic carbon partitioning: its regulation and possibilities for manipulation. Physiol Plant 77: 633–641

Stitt M, Zeeman SC (2012) Starch turnover: Pathways, regulation and role in growth. Curr Opin Plant Biol 15: 282–292

Sulpice R, Flis A, Ivakov AA, Apelt F, Krohn N, Encke B, Abel C, Feil R, Lunn JE, Stitt M (2014) Arabidopsis coordinates the diurnal regulation of carbon allocation and growth across a wide range of Photoperiods. Mol Plant 7: 137–155

Szecowka M, Heise R, Tohge T, Nunes-Nesi A, Vosloh D, Huege J, Feil R, Lunn J, Nikoloski Z, Stitt M, et al (2013) Metabolic Fluxes in an Illuminated Arabidopsis Rosette. Plant Cell 25: 694 LP – 714

Thalmann M, Pazmino D, Seung D, Horrer D, Nigro A, Meier T, Kölling K, Pfeifhofer HW, Zeeman SC, Santelia D (2016) Regulation of Leaf Starch Degradation by Abscisic Acid Is Important for Osmotic Stress Tolerance in Plants. Plant Cell 28: 1860 LP – 1878

Thalmann M, Santelia D (2017) Starch as a determinant of plant fitness under abiotic stress. New Phytol 214: 943–951

Turgeon R (2006) Phloem Loading: How Leaves Gain Their Independence. Bioscience 56: 15

Vanhercke T, Belide S, Taylor MC, El Tahchy A, Okada S, Rolland V, Liu Q, Mitchell M, Shrestha P, Venables I, et al (2019a) Up-regulation of lipid biosynthesis increases the oil content in leaves of Sorghum bicolor. Plant Biotechnol J 17: 220–232

Vanhercke T, Divi UK, El Tahchy A, Liu Q, Mitchell M, Taylor MC, Eastmond PJ, Bryant F, Mechanicos A, Blundell C, et al (2017) Step changes in leaf oil accumulation via iterative metabolic engineering. Metab Eng 39: 237–246

Vanhercke T, Dyer JM, Mullen RT, Kilaru A, Rahman MM, Petrie JR, Green AG, Yurchenko O, Singh SP (2019b) Metabolic engineering for enhanced oil in biomass. Prog Lipid Res 74: 103–129

Vanhercke T, Petrie JR, Singh SP (2014a) Energy densification in vegetative biomass through metabolic engineering. Biocatal Agric Biotechnol 3: 75–80

Vanhercke T, El Tahchy A, Liu Q, Zhou XR, Shrestha P, Divi UK, Ral JP, Mansour MP, Nichols PD, James CN, et al (2014b) Metabolic engineering of biomass for high energy density: Oilseed-like triacylglycerol yields from plant leaves. Plant Biotechnol J 12: 231–239

Vriet C, Welham T, Brachmann A, Marilyn PM, Pike J, Perry J, Parniske M, Sato S, Tabata S, Smith AM, et al (2010) A suite of Lotus japonicus starch mutants reveals both conserved and novel features of starch metabolism. Plant Physiol 154: 643–655

Wagner GJ (1991) Secreting glandular trichomes: More than just hairs. Plant Physiol 96: 675–679

Wagner GJ, Wang E, Shepherd RW (2004) New approaches for studying and exploiting an old protuberance, the plant trichome. Ann Bot 93: 3–11

Weise SE, Liu T, Childs KL, Preiser AL, Katulski HM, Perrin-Porzondek C, Sharkey TD (2019) Transcriptional regulation of the glucose-6-phosphate/phosphate translocator 2 is related to carbon exchange across the chloroplast envelope. Front Plant Sci 10: 827

Weise SE, Schrader SM, Kleinbeck KR, Sharkey TD (2006) Carbon Balance and Circadian Regulation of Hydrolytic and Phosphorolytic Breakdown of Transitory Starch. Plant Physiol 141: 879 LP–886

Weise SE, Weber APM, Sharkey TD (2004) Maltose is the major form of carbon exported from the chloroplast at night. Planta 218: 474–482

Weselake RJ (2016) Engineering oil accumulation in vegetative tissue. Ind. Oil Crop. pp 413–434

Weselake RJ, Taylor DC, Rahman MH, Shah S, Laroche A, McVetty PBE, Harwood JL (2009) Increasing the flow of carbon into seed oil. Biotechnol Adv 27: 866–878

Xu C, Shanklin J (2016) Triacylglycerol Metabolism, Function, and Accumulation in Plant Vegetative Tissues. Annu Rev Plant Biol 67: 179–206

Xu X, Vanhercke T, Shrestha P, Luo J, Akbar S, Konik-Rose C, Venugoban L, Hussain D, Tian L, Singh S, et al (2019) Upregulated Lipid Biosynthesis at the Expense of Starch Production in Potato (Solanum tuberosum) Vegetative Tissues via Simultaneous Downregulation of ADP-Glucose Pyrophosphorylase and Sugar Dependent1 Expressions. Front Plant Sci 10: 1444

Yang Z, Ohlrogge JB (2009) Turnover of fatty acids during natural senescence of arabidopsis, brachypodium, and switchgrass and in arabidopsis β-oxidation mutants. Plant Physiol 150: 1981–1989

Yu L, Fan J, Yan C, Xu C (2018) Starch deficiency enhances lipid biosynthesis and turnover in leaves. Plant Physiol 178: 118–129

Zale J, Jung JH, Kim JY, Pathak B, Karan R, Liu H, Chen X, Wu H, Candreva J, Zhai Z, et al (2016) Metabolic engineering of sugarcane to accumulate energy-dense triacylglycerols in vegetative biomass. Plant Biotechnol J 14: 661–669

Zeeman SC, Kossmann J, Smith AM (2010) Starch: Its Metabolism, Evolution, and Biotechnological Modification in Plants. Annu Rev Plant Biol 61: 209–234

Zhai Z, Liu H, Xu C, Shanklin J (2017) Sugar potentiation of fatty acid and triacylglycerol accumulation. Plant Physiol 175: 696–707

Zhou X-R, Bhandari S, Johnson BS, Kotapati HK, Allen DK, Vanhercke T, Bates PD (2020) Reorganization of Acyl Flux through the Lipid Metabolic Network in Oil-Accumulating Tobacco Leaves. Plant Physiol 182: 739 LP – 755

